# Context-dependent transcriptional regulation by Drosophila Polycomb Response Elements

**DOI:** 10.1101/2022.05.11.491498

**Authors:** Rory T. Coleman, Gary Struhl

**Author notes:** Present address: The Rockefeller University,1230 York Avenue, New York, N.Y. 10065.

## Abstract

Polycomb Response Elements (PREs) are *cis*-acting DNA sequences that confer heritable states of Drosophila HOX gene expression by anchoring Polycomb and Trithorax Group (PcG and TrxG) chromatin modifiers. PREs are also associated with hundreds of other Drosophila genes, most of which are regulated dynamically in response to developmental and physiological context, rather than heritably like HOX genes. Here, we assess the role(s) PREs play at these other loci by analyzing how genomic inserts of a transgenic form of the HOX gene *Ultrabithorax* (*Ubx*) can both control and respond to neighboring genes depending on the presence of a single, excisable PRE. Our results support the view that PREs and their associated PcG and TrxG modifiers act primarily to confer quantitative, rather than qualitative, influences on gene expression with the response of any given gene depending on how it integrates this information with other regulatory elements in the local genomic milieu. They also show that PREs can act on neighboring genes selectively and at remarkably long range, but that any given gene can be susceptible or impervious to PRE/PcG/TrxG input depending on context. Finally, we find that transcription and PRE/PcG-dependent silencing are not mutually exclusive: a *Ubx* transgene inside the intron of a continuously transcribed “host” gene is nevertheless silenced by its resident PRE. We posit that the widely accepted roles of PcG and TrxG complexes in maintaining heritable states of gene expression apply only to a limited coterie of target genes such as HOX genes that are evolutionarily selected to exclude regulatory elements that can over-ride this control.

## Introduction

During development, groups of cells commit to forming particular tissues or body parts by heritably activating some control genes whilst silencing others. This phenomenon is epitomized by the specification of body segments by HOX genes in most animals and depends on chromatin modifying enzyme complexes that are highly conserved in animals and plants. These include Polycomb Group (PcG) complexes, which maintain HOX gene silencing, and Trithorax Group (TrxG) complexes, which maintain HOX gene activity.

In Drosophila, both the activation and heritable silencing of HOX genes depends on *cis*-acting regulatory elements, known as Polycomb Response Elements (PREs) (Simon et al., 1993; Christen and Bienz, 1994; Chan et al., 1994), which act by recruiting PcG and TrxG complexes [reviewed in (Müller and Kassis, 2006; Bauer et al., 2016; Erokhin et al., 2018)]. Molecular studies of HOX genes have provided a paradigm for how such complexes maintain heritable states of gene expression.

Early in development, HOX genes that are initially repressed become substrates for two classes of PcG complexes: Polycomb Repressive Complexes 1 and 2 (PRC1 and PRC2). PRC2 contains a methyltransferase, Enhancer of Zeste [E(z)], that catalyzes trimethylation of histone H3 at lysine 27 (H3K27me3)—a repressive epigenetic mark that is inherited and copied in descendant cells after each cycle of DNA replication (Czermin et al., 2002; Müller et al., 2002; Kuzmichev et al., 2002; Cao et al., 2002; Hansen et al., 2008; Margueron et al., 2009; Pengelly et al., 2013; Coleman and Struhl, 2017; Laprell et al., 2017; Reverón-Gómez et al., 2018; Poepsel et al., 2018). PRC1 then binds to H3K27me3 and further modifies chromatin both to confer transcriptional repression and to reinforce recruitment of PRC2 (Cao et al., 2002; Min et al., 2003; Fischle et al., 2003; Wang et al., 2004a; Kahn et al., 2016; Blackledge et al., 2020; Dobrinić et al., 2021).

By contrast, genes that are initially activated become substrates for TrxG complexes. These take up residence at the promoter and catalyze chromatin modifications such as H3K4 and H3K36 methylation that allow gene expression by blocking PRE/PRC2-dependent H3K27 trimethylation of the structural gene (Klymenko and Müller, 2004; Papp and Müller, 2006; Tie et al., 2009; Schuettengruber et al., 2009; Yuan et al., 2011; Schmitges et al., 2011; Tie et al., 2014; Streubel et al., 2018; Finogenova et al., 2020). Loss of TrxG activity results in inappropriate H3K27 trimethylation and silencing of HOX genes (Klymenko and Müller, 2004; Papp and Müller, 2006; Tie et al., 2009). This phenotype is reversed by simultaneous removal of PRC2, which results in the loss of HOX gene silencing even in the absence of TrxG function (Klymenko and Müller, 2004). Thus, HOX gene silencing can be viewed as a default state that is established and maintained by PRE/PRC activity in the absence of TrxG complex engagement (Müller and Kassis, 2006).

Although PREs were first defined by their roles in heritable HOX gene expression, it is now apparent that they are widely dispersed throughout the genome and regulate hundreds if not thousands of *Drosophila* genes, most of which are not expressed in a heritable fashion (Schwartz et al., 2006; Oktaba et al., 2008; Kuroda et al., 2020). Moreover, many such PRE regulated genes in *Drosophila* have homologues that are subject to PcG regulation in vertebrates (Boyer et al., 2006; Bracken et al., 2006), although vertebrates depend primarily on means other than PREs to anchor PRCs to their targets [reviewed in (Bauer et al., 2016; Yu et al., 2019; Blackledge and Klose, 2021)].

These observations challenge the view that the primary role of PcG and TrxG complexes is to confer all-or-none, heritable states of gene expression [reviewed in (Schwartz and Pirrotta, 2007; Sawarkar and Paro, 2010)]. On the other hand, segment determination depends critically on the capacity of cells to choose the appropriate code of HOX gene activities and then maintain that choice in all of their descendants (Lewis, 1978; Struhl, 1981; Duncan, 1982; Struhl and Akam, 1985; Beuchle et al., 2001). This poses the question of how the epigenetic roles ascribed to these complexes in the case of HOX genes relates to their regulation of other target genes that are expressed more dynamically in response to developmental and physiological context.

Here, we address this question by assaying random genomic insertions of a *lacZ* reporter form of the classical Drosophila HOX gene *Ultrabithorax* (*Ubx*) that depends on an excisable PRE within a Flp-out cassette (Coleman and Struhl, 2017). Previously, we used one such *>PRE>Ubx.lacZ* transgene insertion to establish that the PRE can maintain heritable silencing by recruiting PRC2 to copy the H3K27me3 epigenetic “mark” following DNA replication (Coleman and Struhl, 2017). Here, we focus on other *>PRE>Ubx.lacZ* insertions that reveal the potential for remarkably diverse, context-dependent properties of transcriptional regulation mediated by the transgene PRE.

These include (i) the capacity of the PRE to exert a quantitative, rather than a qualitative, control on target gene transcription, (ii) the ability of the PRE to impose silencing on previously active genes or to sustain the activity of target genes that would otherwise cease to be expressed, (iii) a critical role of genomic and developmental context in rendering a promoter susceptible or refractory to PRE mediated regulation, (iv) the capacity of the PRE to act in cis and at long range on select target genes without influencing the expression of intervening genes, and (v) the ability of the PRE to maintain silencing of the *Ubx* promoter even when the entire transgene resides in the intron of a “host” gene that is continuously expressed.

Based on these results, we view PRE’s as pervasive, multi-functional regulatory elements that act via PcG and TrxG complexes to provide repressive and activating inputs, with the outcome at any given promoter depending on how this information is integrated with the regulatory activities of other cis-acting elements. From this perspective, we posit that the canonical roles of PcG and TrxG complexes in conferring heritable states of gene expression do not reflect their fundamental role, but rather an attribute of the target genes themselves— in the case of HOX genes, the biological imperative to function as determinants of segmental state. Accordingly, genes that are heritably regulated by PREs may be under stringent evolutionary selection to exploit chromatin modifications as epigenetic marks and to exclude other cis-acting regulatory elements that can over-ride the ON or OFF states they confer.

## Results

### Probing context-dependent PRE function using a *>PRE>UZ* transgene

Upon fertilization, *Drosophila* embryos undergo a series of syncytial nuclear divisions during which the nuclei migrate to the egg periphery and are incorporated into cells. The resulting blastoderm is then partitioned into distinct head, thoracic and abdominal segmental primordia via the heritable activation of different combinations of HOX genes in each primordium. The classic HOX gene *Ultrabithorax* (*Ubx*) is activated in cells that will give rise to parasegments 5-13, corresponding approximately to the third thoracic and first eight abdominal segments (Lewis, 1978; Struhl, 1984). *Ubx* expression can be recapitulated by *Ubx.lacZ* (*UZ*) reporter transgenes composed of four elements from the native gene: an early embryo enhancer (EE), a later acting imaginal disc enhancer (DE), a Polycomb Response Element (PRE), and the *Ubx* promoter driving transcription of the βGalactose coding sequence [(Chan et al., 1994; Christen and Bienz, 1994; Bienz and Müller, 1995; Pirrotta et al., 1995); reviewed in (Schwartz and Pirrotta, 2007)]. Here, we use one such transgene, *>PRE>UZ*, in which the PRE is embedded inside a Flp-out cassette (Struhl and Basler, 1993), allowing it to be excised by recombination between Flp-recombinase target sites (*>*) at each end [Fig. 1A; Experimental Methods; Table 1; (Coleman and Struhl, 2017)]. Importantly, the *>PRE>* cassette also contains a *Tubulin1α.CD2* (*Tub.CD2*) mini-gene, which expresses the rat CD2 protein in all cells under the control of the *Tub* promoter (Jiang and Struhl, 1995). As a consequence, all of the clonal descendants of cells in which excision occurs can be marked by the absence of CD2.

**Figure 1.**
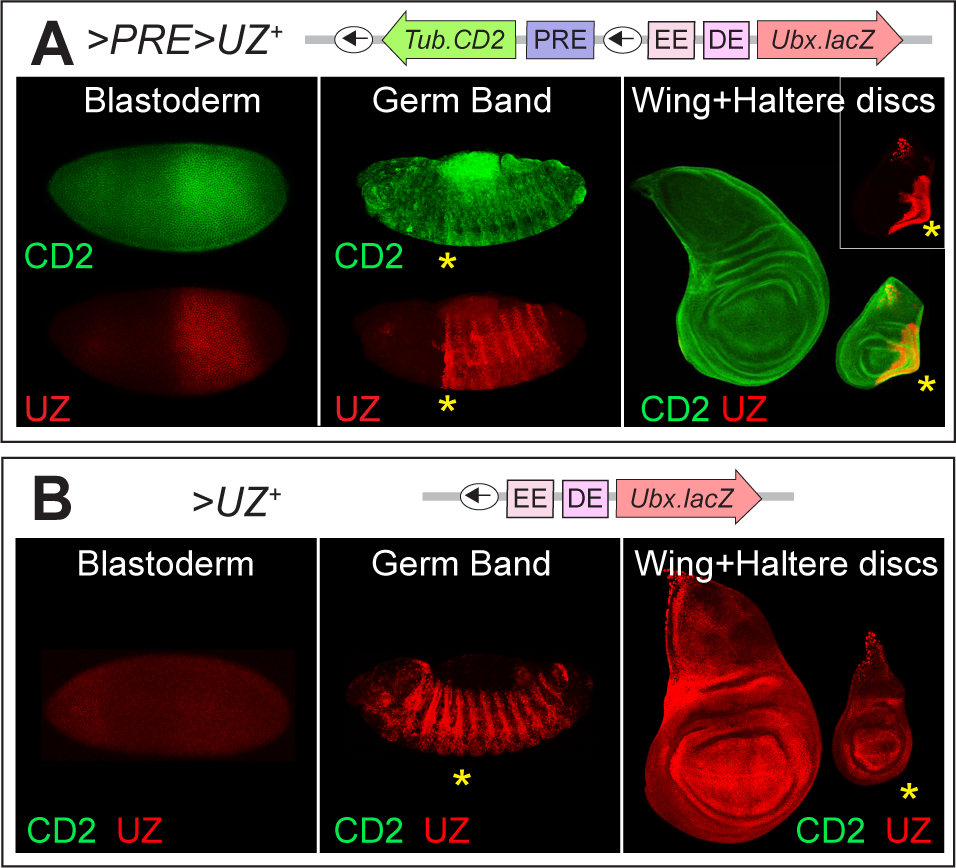
Structure and activity of the canonical *>PRE>UZ^+^* transgene with and without the *>PRE>* cassette. **A)** The intact *>PRE>UZ^+^* transgene contains an *Ubx.lacZ* mini-gene composed of the *Ubx* promoter fused to the lacZ coding sequence (*UZ*) positioned downstream of three well characterized elements from the native locus: an early enhancer (EE), an imaginal disc enhancer (DE) and the classic 1.6kb *bxd* PRE element embedded within a “Flp-out” cassette flanked by Flp Recombinase Targets (FRT’s, small arrows) [Experimental Methods; Table 1; (Coleman and Struhl, 2017)]. The cassette also contains an upstream *Tub.CD2* mini-gene that points away from the *UZ* mini-gene and normally expresses rat CD2 protein uniformly throughout development, allowing cells carrying the *>PRE>* to be identified by staining for CD2. The *>PRE>UZ^+^* transgene also includes an ∼ 8kb rescuing genomic fragment from the *yellow* (*y*) gene downstream from the *UZ* mini-gene, which serves as a genetic marker (not shown). Expression of CD2 (green) and the *>PRE>UZ^+^* transgene (the latter monitored by βGal staining, red, referred to subsequently as UZ) in blastoderm embryos, shortened germ band embryos and in the large wing and smaller haltere imaginal discs from a late 3^rd^ larva as indicated (here and in subsequent figures, anterior is to the left and dorsal to the top; and nuclei counterstained as needed with DAPI, blue). UZ is initially activated like native Ubx in a broad domain posterior domain during the blastoderm stage which quickly sharpens to parasegments 6-12 after the onset of gastrulation, as apparent in the germ band shortened embryos (note that native Ubx is expressed similarly although in a larger domain encompassing parasegments 5-13). The sharp border at the anterior boundary of parasegment 6, subdivides the third thoracic segment into distinct A and P compartments, all of whose descendants inherit their initial off and on states of UZ expression for the rest of development: UZ is off in the imaginal discs that will form the wing (second thoracic) and haltere (third thoracic) appendages of the adult, except in the P compartment of the haltere disc, where it is expressed in all cells (here and in all subsequent figures, the P compartment is marked by a yellow asterisk). CD2 is expressed ubiquitously from the blastoderm stage onwards, albeit up-regulated transiently with early UZ expression in the blastoderm. This up-regulation declines during embryogenesis, resulting in uniform, ubiquitous expression in all the imaginal discs. **B)** The PRE-excised form of the *>PRE>UZ^+^*transgene *(>UZ^+^*) is not initially activated during the blastoderm stage, but instead comes on later, in all segments, as visualized in the germ band shortened embryo and in the imaginal discs (here and in Figs. 2-5, the *>UZ* genotype was confirmed by absence of CD2 expression). Hence, the PRE plays an essential early role, like that of the EE, in the initial activation of the *UZ^+^* promoter in parasegments 6-12. Conversely, it is required to maintain the silenced state in the remaining parasegments during subsequent development.

**Table 1.**
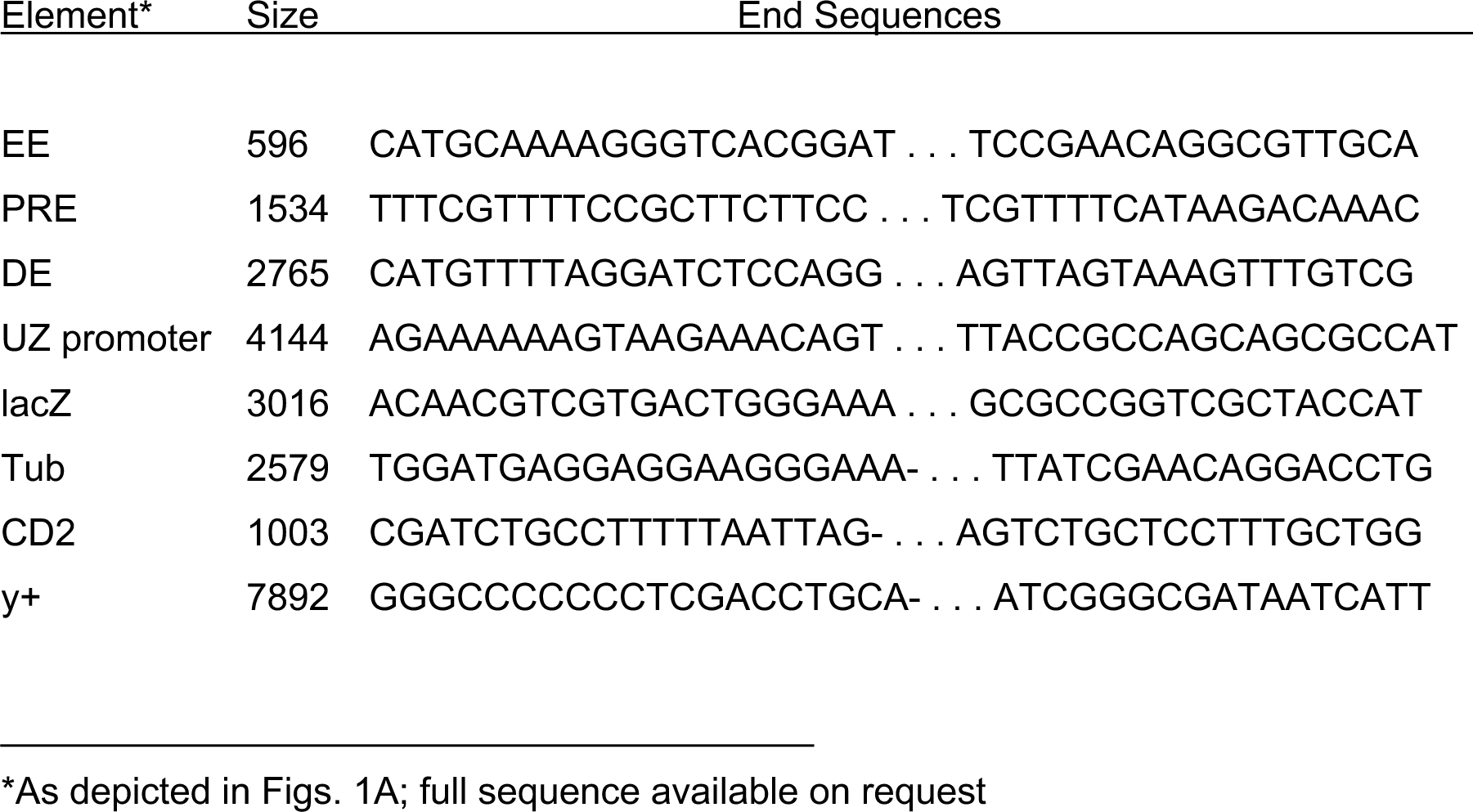
Composition of the *>PRE>UZ* transgene.

We have analyzed seven genomic insertions of the *>PRE>UZ* transgene (Table 2), of which four show the expected *Ubx*-like pattern of expression throughout development [(Coleman and Struhl, 2017); Fig. 1A]. However, the remaining three show different patterns that result from interactions with neighboring genomic DNA. Here, we present a detailed analysis of these transgenes to gain insight into how the regulatory inputs of PREs and their anchored chromatin modifiers are integrated with those of other cis-acting elements and their associated factors.

**Table 2.** Genomic insertion sites of the *>PRE>UZ* transgenes.

| Transgene |  | Insertion site* |
| --- | --- | --- |
| >PRE>UZ<sup>+</sup> | 55B11** | 2R:18171538/9 |
| >PRE>UZ<sup>Hmx</sup> | 90B4 | 3R:17547893/4 |
| >PRE>UZ<sup>Wnt</sup> | 27F2 | 2L:7356049/50 |
| >PRE>UZ<sup>pum</sup> | 85C11 | 3R:9158098/9 |

\* Flybase release r6.44; orientation as in Figs. 1,2,3 and 5.
\*\* Three other >PRE>UZ transgene insertions, located at 59B2, 82B1, and 85A10 behave similarly to the canonical >PRE>UZ<sup>+</sup> transgene located at 55B11.

To provide a basis for comparison, we first present the properties of a canonical, “well-behaved” transgene inserted at chromosomal location 55C [henceforth *>PRE>UZ^+^*; (Coleman and Struhl, 2017)]. The *>PRE>UZ^+^* transgene generates a pattern of β-Galactosidase (henceforth UZ) expression similar to that of native Ubx, except limited to parasegments 6-12 rather than 5-13 [likely because the EE fragment lacks the parasegment 5 specific, *anterobithorax* enhancer element (Peifer and Bender, 1986)].

UZ expression is first detected at the cellular blastoderm stage in a broad domain that corresponds to the founding cells of parasegments 6-12 (Fig. 1A, left panel). Beginning with the onset of gastrulation, expression rises rapidly to peak levels in these parasegments, with a sharp border coinciding with the anterior boundary of parasegment 6 (Fig. 1A, middle panel; here and in subsequent figures, UZ expression is shown in red, CD2 expression in green, and parasegment 6 derivatives are indicated by a yellow asterisk). This boundary subdivides the third thoracic segment of the embryo, including the primordia destined to give rise to haltere and third leg imaginal discs of the larva, into anterior (A) and posterior (P) compartments belonging, respectively, to parasegments 5 and 6. The ON and OFF states of UZ expression are then inherited for the remainder of development, UZ being present in the P, but not the A, compartments of the haltere and third leg discs, and absent in all the more anterior thoracic and head discs, including the wing disc (Fig. 1A, right panel).

Importantly, the *Tub.CD2* gene within the *>PRE>* cassette is expressed ubiquitously throughout development even though the PRE abuts the *Tub* promoter within the cassette. Hence, in this context, the *Tub* promoter appears refractory to the presence of the PRE. The *Tub* promoter does, however, respond transiently to the EE on the other side of the PRE, as its expression is initially up-regulated in the same broad domain as UZ in early embryos (Fig. 1A, left panel).

In contrast, the *>UZ^+^*transgene generated by germ-line excision of the *>PRE>* cassette is ubiquitously expressed in embryos as well as larvae, including in all of the thoracic discs (Fig. 1B, middle and right panels), indicating that the PRE is required for the maintenance of the OFF state. However, unlike the intact *>PRE>UZ^+^* transgene, the *>UZ^+^* transgene is not activated during the blastoderm stage (Fig. 1B, left panel). Instead, its expression is first detected only later, in extended germ band embryos, when it is derepressed in all segments. Thus, the PRE plays an early activating role that is equivalent to that of the EE in parasegments 6-12 (Coleman and Struhl, 2017), in addition to its repressive role in silencing UZ expression in the rest of the body. This is consistent with previous work suggesting that PREs constitutively anchor both TrxG and PcG complexes, which then exert their activating or repressing input at the promoter as dictated by the initial activated or repressed state of the EE (Christen and Bienz, 1994; Chan et al., 1994; Klymenko and Müller, 2004; Papp and Müller, 2006; Kahn et al., 2006).

Below, we analyze three *>PRE>Ubx* insertions that behave differently from the canonical *>PRE>UZ^+^* insertion, using the ability to excise the PRE to probe its impact on the *UZ* and *Tub.CD2* minigenes within the transgene as well as on neighboring genes.

### Regulation of a *>PRE>UZ* transgene by a putative PRE in neighboring genomic DNA

The first atypical insertion, *>PRE>UZ^Hmx^*, is located close to the *H6-like-homeobox* (*Hmx*) gene (Fig. 2A; Table 2). This transgene behaves similarly to the canonical *>PRE>UZ^+^* transgene throughout development, except that the *Tub.CD2* mini-gene within the *>PRE>* cassette is strongly repressed in all of the thoracic discs, except in the P compartments of the haltere and third leg discs where it coincides with UZ expression (Fig. 2B; right panel). However, in contrast to the *>PRE>UZ^+^*transgene, excision of the PRE, whether during development or in the germ-line of a parent, does not result in a loss of silencing (Fig. 2C). Instead, the resulting *>UZ^Hmx^*transgene is expressed like the intact *>PRE>UZ^Hmx^* transgene, being activated in early embryos in parasegments 6-12, but repressed in all the remaining parasegments thereafter (Fig. 2C).

**Figure 2.**
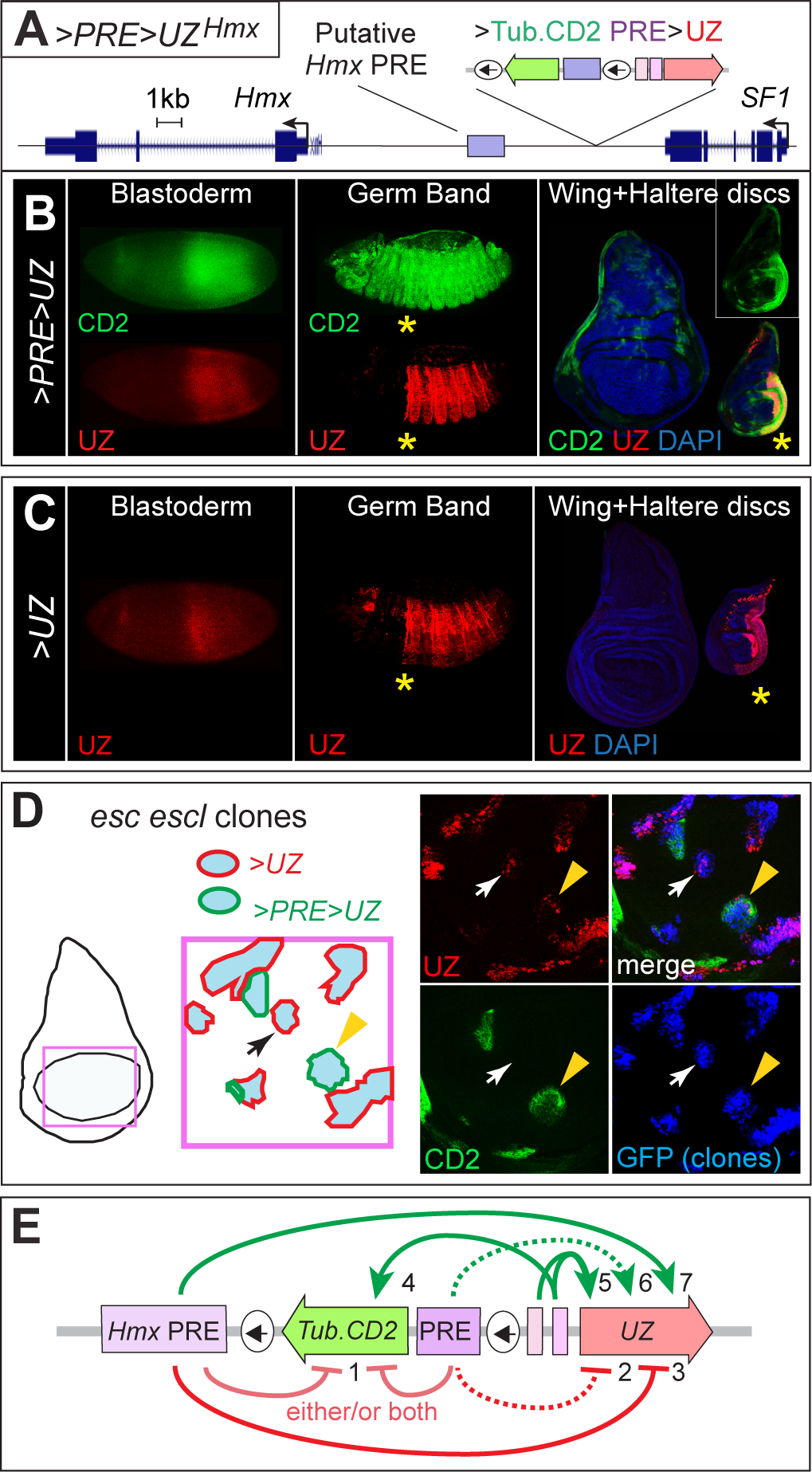
Redundant silencing of the *>PRE>UZ^Hmx^* transgene insert close to the *Hmx* locus by a putative PRE in neighboring genomic DNA. **A)** Genomic site of the *>PRE>UZ^Hmx^*transgene inserted near the *Hmx* locus (the orientation of the *Tub.CD2*, *>PRE>* and *UZ* components of the transgene are indicated above the insertion site; not to scale). A putative PRE, assigned on the basis of ChIP- chip data (Schuettengruber et al., 2009), is shown in purple. **B)** CD2 and UZ expression from the *>PRE>UZ^Hmx^*transgene in blastoderm, germ band shortened embryos and in wing and haltere discs (as in Fig. 1). UZ and CD2 expression in embryos are indistinguishable from their expression in *>PRE>UZ^+^*embryos (Fig. 1), but CD2 is silenced in most cells of the wing and haltere discs, with the exception of the P compartment of the haltere disc, where UZ is expressed. **C)** UZ expression in *>UZ^Hmx^* embryos and imaginal discs (as in B). In contrast to the *>PRE>UZ^+^* transgene (Fig. 1B), UZ expression is activated normally at the cellular blastoderm stage and then maintained in the appropriate “on” and “off” states thereafter (except for a patch of UZ expressing cells in the head of the embryo), despite the absence of the transgene PRE. **D)** Heritable silencing of the *>PRE>UZ^Hmx^*and *>UZ^Hmx^* transgenes depends on PRC2. The left panel depicts a wing disc containing clones of homozygous *esc escl* double mutant cells (labeled positively by GFP staining; blue shading) in the prospective wing (purple box) that carry either the intact or the PRE-excised form of the *>PRE>UZ^Hmx^* transgene (green or red outlines respectively). The right panel shows the independent stains for UZ, CD2 and GFP, as well as the merge. An *esc^-^ escl^-^* mutant clone carrying the *>PRE>UZ^Hmx^* transgene is indicated by a yellow arrowhead (note the co-expression of both CD2 and UZ); another clone, which we infer carries the *>UZ^Hmx^* transgene, is indicated by a white arrow (expresses UZ, but not CD2). **E)** Diagram of tested and inferred interactions between the PREs, enhancers and promoters associated with the *>PRE>UZ^Wnt^*transgene. We infer the presence of a putative “Hmx PRE” that acts either alone or together with the PRE in the *>PRE>UZ^Hmx^* transgene to silence both the *Tub* (1,2) and *UZ* (3) promoters, as well as to initiate expression of the UZ promoter (6,7). The EE drives early expression of both the *Tub* (4) and *UZ* (5) promoters. Silencing and activation of the UZ promoter by the transgene PRE are shown as dotted lines (2,6) because they cannot be confirmed by PRE excision due to the redundant action of the inferred Hmx PRE.

A simple explanation for the heritable silencing of the *>UZ^Hmx^* transgene is that absence of the transgene PRE is compensated for by a PRE in neighboring genomic DNA (Figs. 2A). If so, silencing of the excised *>UZ^Hmx^* transgene should depend on PRC2 activity. To test this, we used Flp-mediated mitotic recombination to generate clones of cells that lack both Extra Sex Combs (Esc) and its paralogue Esc-like (Escl), which semi-redundantly provide an essential component of PRC2 (Struhl, 1981; Ohno et al., 2008; Margueron et al., 2009), in animals that carry the *>PRE>UZ^Hmx^*transgene.

Because Flp catalyzes both *>PRE>* cassette excision as well as FRT-mediated mitotic recombination, some of the resulting *esc^−^ escl*^−^ clones retain the intact *>PRE>UZ^Hmx^* transgene, whereas others carry the PRE-excised, *>UZ^Hmx^* form (Experimental Methods). In wing discs, in which both CD2 and UZ would otherwise be silenced, we observe two classes of *esc^−^ escl*^−^ clones: those that express only UZ and those that express both UZ and CD2 (Fig. 2D). Hence, we infer that silencing of the *>UZ^Hmx^* transgene depends on a redundant PRE located in neighboring genomic DNA, and silencing of the *Tub* promoter in the *>PRE>UZ^Hmx^* transgene depends on either (or both) the transgene and genomic PREs (Fig. 2E).

Thus, it appears that at least some, and possibly many, of the hundreds of putative PREs scattered throughout the genome (Schwartz et al., 2006; Kwong et al., 2008; Oktaba et al., 2008; Bredesen and Rehmsmeier, 2019) can substitute functionally for the PRE within the *>PRE>UZ* transgene. This includes providing both the early role in activating the *UZ* promoter in parasegments 6-12 (Fig. 2C, left panel) as well as the silencing role in repressing the promoter in the remaining parasegments (Fig. 2C, right panel). Strikingly, the *Tub* promoter within the intact *>PRE>UZ^Hmx^* transgene is also silenced by PRC2 activity (Fig. 2D), even though it is impervious to repression by the abutting PRE in the canonical *>PRE>UZ^+^*transgene (Fig. 2A). This result establishes that a given promotor can be either refractory or susceptible to PRE/PRC-dependent silencing in a manner that depends on genomic context rather than an intrinsic property of the promoter (see below and Discussion).

### Context-dependent integration of enhancer and PRE inputs revealed by a ***>PRE>UZ* transgene in the major *Wnt* complex.**

The second, atypical insertion, *>PRE>UZ^Wnt^*, is located in a gene complex that contains four of the seven Drosophila *Wnt* genes (*Wnt4*, *wg*, *Wnt6* and *Wnt10*; Fig. 3A). *Wnt4*, *wg*, and *Wnt6* are expressed similarly in the imaginal discs (Gieseler et al., 1999; Janson et al., 2001), suggesting that they are coordinately regulated by some of the same cis-acting regulatory elements. However *Wnt6* expression in embryos differs from *Wnt4* and *wg* in being expressed only weakly and late, in the gut, and *Wnt10* is expressed weakly, or not at all, in both embryos and the imaginal discs (Gieseler et al., 1999; Janson et al., 2001). As we describe below, the behavior of the *>PRE>UZ^Wnt^*transgene reveals an unexpectedly complex and context-dependent integration of available enhancer and PRE elements by both the *UZ* and *Tub* promoters within the transgene as well as by the promoter of the endogenous *wg* gene located ∼50 kb away.

**Figure 3.**
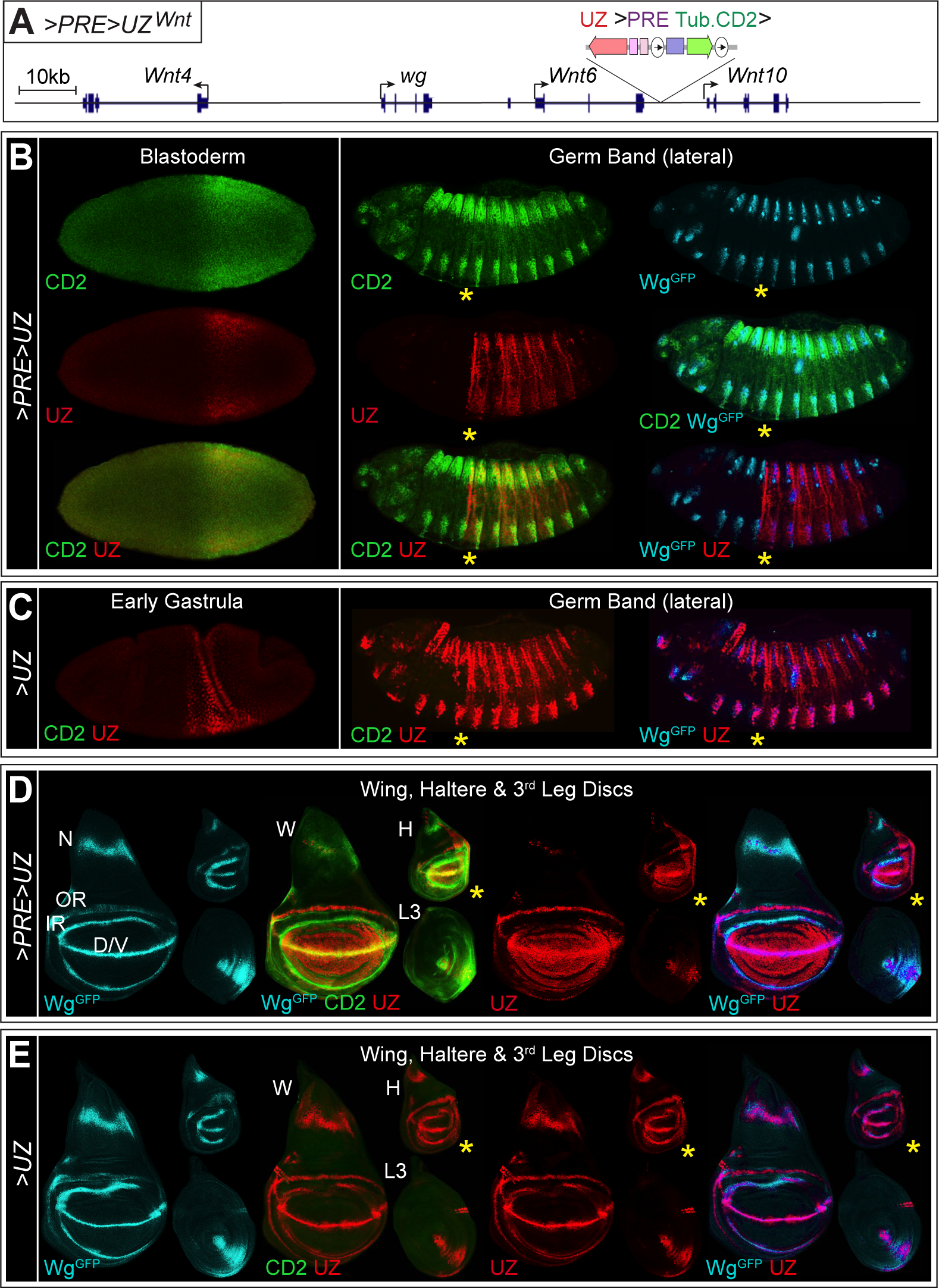
Integration of enhancer and PRE inputs regulating a *>PRE>UZ* transgene in the *Wnt* complex. **A)** Genomic site and orientation of the *>PRE>UZ^Wnt^*transgene in the main Wnt gene complex containing *Wnt4*, *wg*, *Wnt6* and *Wnt10* (as in Fig. 2; the transgenes is not to scale). **B)** UZ, CD2 and Wg expression in blastoderm and germ band shortened *>PRE>UZ^Wnt^* embryos (oriented and annotated as in Fig. 2). Both CD2 and Wg expression are usually monitored using mouse monoclonal antisera; hence, to assay their co-expression, we used embryos that carry the *>PRE>UZ^Wnt^* transgene in trans to a *wg^GFP^* knock-in allele that is normally expressed and fully functional (Port et al., 2014) and stained for GFP (turquoise). UZ expression is initially activated in parasegments 6-12 and silenced in the remaining parasegments as observed for the *>PRE>UZ^+^*and *>PRE>UZ^Hmx^* transgenes (Figs. 1A, 2B). However, CD2 appears coincident with that of Wg^GFP^ after the onset of gastrulation, indicating that the *Tub* promoter, but not the *UZ* promoter, has been coopted by Wnt complex embryonic enhancers and silencers. **C)** UZ and Wg^GFP^ expression in early gastrula and germ band shortened *>UZ^Wnt^* embryos. In contrast to the canonical *>UZ^+^* transgene (Fig. 1B), but similar to the *>UZ^Hmx^* transgene (Fig. 2C), UZ expression is activated in parasegments 6-12 (albeit slightly later, at the onset of gastrulation instead of during the blastoderm stage) and silenced in the remaining parasegments, with the notable exception of cells that express wg^GFP^. Hence, we infer (i) the existence of a redundant PRE in the *wg*/*Wnt* complex that can provide both the early activation and subsequent silencing functions that would otherwise depend on the excised transgene PRE, and (ii) that in the absence of the transgene PRE, Wnt complex enhancers over-ride the silencing activity of this putative Wnt PRE to drive UZ expression in cells that express Wg^GFP^. At least one PRE implicated in Wg-dependent regeneration has been identified, **D)** UZ, CD2 and Wg^GFP^ expression in mature wing (W), haltere (H), and third leg (L3) discs of *>PRE>UZ^Wnt^* larvae. Left-most image: Wg is normally expressed in the wing disc in a broad stripe in the prospective notum (N), inner and outer rings in the prospective wing hinge (IR, OR), and in cells flanking the D/V compartment boundary (D/V) in the prospective wing blade (see also Fig. 4A), and is expressed similarly in the haltere disc, except that no D/V or IR expression is observed in the P compartment; Wg is also expressed in a ventral wedge in the A compartment of the third leg disc, abutting the A/P compartment boundary. Remaining images: As in the embryo, CD2 expression appears similar if not identical to that of Wg^GFP^ in all three discs, except being greatly reduced or absent in the OR in the wing and haltere discs (see also Fig. 4A). However, in contrast to the embryo, UZ is now expressed in a wg-like pattern in all three discs, super- imposed on the normal pattern of UZ expression observed for the canonical *>PRE>UZ^+^*transgene (normally restricted to the P compartments of the haltere and third leg discs). Hence, imaginal disc Wnt enhancers appear to over-ride silencing of the UZ promoter that would otherwise be maintained by the transgene PRE. However, the pattern of UZ expression departs from that of Wg in three respects. First, the stripe of UZ expression in the notum is abnormally narrow; second, UZ is not expressed in the IR in either the haltere or wing disc (these cells appear turquoise, rather than magenta in the Wg^GFP^ UZ merged image), and third, UZ is expressed throughout the prospective wing blade (appears red in the CD2, UZ and Wg^GFP^ UZ merged images) rather than being restricted to the cells flanking the D/V compartment boundary (shown in more detail in Fig. 4A). **E)** Expression of UZ and Wg^GFP^ expression in wing, haltere, and third leg discs of *>UZ^Wnt^*larvae. In the absence of the transgene PRE, UZ expression now appears very similar to both Wg^GFP^, as well as CD2 from the intact *>PRE>UZ^Wnt^* transgene (**D**), *e.g.*, showing the normal broad stripe in the notum, partial restoration of expression in the IR of the hinge, and tight restriction of expression to D/V border cells in the wing. However, a notable exception is the pattern of UZ, which continues to be expressed in the OR of *>UZ^Wnt^* wing discs, in contrast to CD2 which is repressed in the OR of *>PRE>UZ^Wnt^* discs (see Figs. 4A-C).

In embryos, the expression of UZ generated by the intact *>PRE>UZ^Wnt^* transgene (Fig. 3B) appears identical to that of the canonical *>PRE>UZ^+^* transgene (Fig. 1B).

However, this is not the case for the *Tub.CD2* mini-gene inside the >PRE> cassette. Instead, following the onset of gastrulation, CD2 expression deviates from its characteristic early pattern (Figs. 1A and 3B) and is up-regulated in a pattern similar to that of *wg*, most notably showing peak expression in a thin stripe of cells just anterior to each parasegment boundary, coincident with the stripes of native Wg expression, whilst being repressed elsewhere [Fig. 3B, right panel; Wg expression is monitored here and in other panels except Fig. 4C by visualizing GFP encoded by a fully functional *wg^GFP^* knock-in allele (Port et al., 2014); Experimental Methods]. Thus, it appears that the *Tub* promoter is now responding to regulatory elements that normally control the *wg* and *Wnt4* promoters, even though the neighboring *UZ* promoter is refractory to their influence.

**Figure 4.**
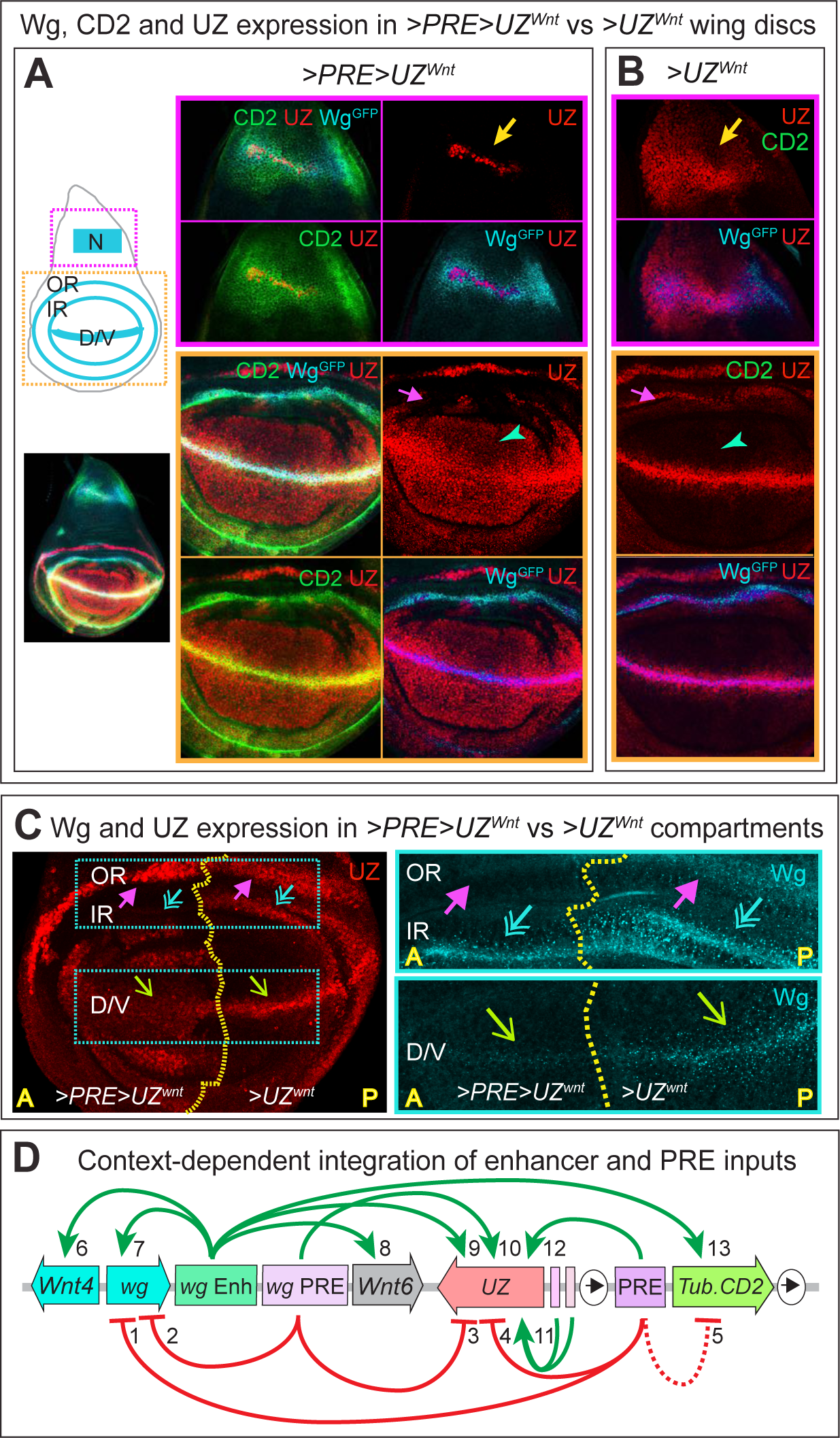
Integration of PRE and enhancer inputs by the *UZ*, *Tub* and native *wg* promoters associated with the *>PRE>UZ^Wnt^* transgene. **A,B)** CD2, Wg^GFP^ and UZ expression of wing discs heterozygous for the intact (**A**) or PRE-excised (**B**) forms of the *>PRE>UZ^Wnt^*transgene. An annotated image of a *>PRE>UZ^Wnt^* wing disc is shown on the left, with the regions shown at higher magnification to the right boxed in purple (prospective notum; top panels) or orange (prospective wing hinge and wing blade; bottom panels). In the prospective notum of the *>PRE>UZ^Wnt^* wing disc (**A**), CD2 expression is similar to, but somewhat broader than, Wg^GFP^ expression, whereas UZ expression is restricted to a narrow stripe that appears to coincide with cells that express peak levels of Wg^GFP^ and CD2 (yellow arrow). However, in the *>UZ^Wnt^* wing disc (**B**), UZ expression is now broad, mimicking that of CD2 observed for the intact transgene (yellow arrow; note that for both genotypes, UZ is expressed only in a central portion of the notum stripe, and not in the portion of the stripe that extends to the posterior edge of the disc, possibly because these two portions of the stripe are under the control of distinct constellations of regulatory factors). In the prospective wing hinge and wing blade, UZ expressed from the intact *>PRE>UZ^Wnt^*transgene (**A**) appears coincident with Wg^GFP^ in the outer ring (OR, depicted in the annotated image to the left; see also Fig. 3D), but is absent from the IR (pink arrow in the UZ only image; the IR appears green and turquoise, respectively, in the CD2 Wg^GFP^ UZ and Wg^GFP^ UZ merged images, rather than yellow and magenta as observed for coincident expression of all three signals along the D/V border in the prospective wing). Conversely, CD2 expression is absent from the OR, which appears red, rather than yellow in the CD2 UZ merge and turquoise in the Wg^GFP^ UZ merge (see also **D**). However, in the absence of the PRE (**B**), UZ is now expressed in the IR, coincidentally with Wg^GFP^ (appears magenta in the Wg^GFP^ UZ merge). Finally, UZ is expressed throughout the prospective wing blade primordium in the *>PRE>UZ^Wnt^* wing disc (**A**; green arrowhead), but restricted to cells flanking the D/V boundary in the *>UZ^Wnt^*wing disc (**B**). **C)** UZ and Wg expression in a wing disc homozygous for the *>PRE>UZ^Wnt^* transgene that retains the PRE in A compartment cells, but lacks the PRE in P compartment cells owing to P-compartment specific expression of Flp recombinase under *hh.Gal4/UAS* control. Left panel: UZ is expressed in OR cells in both the A and P compartments (magenta arrows), but only in IR cells in the P compartment (turquoise double arrows), indicating that the OR, but not the IR, *Wnt* enhancers can over-ride silencing by the transgene PRE. As in (**A**), the presence of the PRE in A compartment cells also sustains UZ expression throughout the prospective wing blade, which would otherwise be restricted to D/V border cells, as it is in P compartment cells (green arrows). Right panel: the presence of the PRE in A cells also abolishes native Wg expression in OR cells, in contrast to the P compartment where OR cells express Wg (purple arrows), a deficiency confirmed by the loss of a specific wing hinge structure, the axillary cord, specified by Wg expressed by OR cells (Neumann and Cohen, 1996); it also reduces the level of Wg expression in IR and D/V border cells (compare expression levels in A and P (turquoise double arrows and green arrows, respectively). **D)** Diagram of tested and inferred interactions between the PREs, enhancers and promoters associated with the *>PRE>UZ^Wnt^*transgene. The PRE in the *>PRE>UZ^Wnt^* transgene silences or down-regulates the native *wg* promoter (1), and possibly the *Tub* promoter (5; dashed line) in OR, IR and D/V border cells. It also silences the *UZ* promoter in the embryo and IR cells (4), restricts the domain of UZ expression in notum cells (4), and sustains the activity of the *UZ* promoter in presumptive wing tissue of the imaginal disc (12). PRE(s) in the Wnt complex repress damage induced expression of the native *wg* promoter in the imaginal discs (Harris et al., 2016) (2) and also appear to regulate the *UZ* promoter in the embryo, activating it in parasegments 6-12 (10) and silencing it elsewhere (3; for simplicity, all Wnt PREs are depicted as a single element). Enhancers in the Wnt complex activate *wg* (7) and as well as *Wnt4* and *Wnt6* (6,8), the *Tub.CD2* mini-gene within the transgene in most *wg* expressing cells (13), and the *UZ* gene in some or most *wg* expressing cells, depending on the presence or absence of the transgene PRE (9). The EE and DE enhancers drive expression of UZ promoter, respectively, in the embryo and posterior compartment of the haltere and third leg disc (11). The Wnt enhancers and transgene PRE also regulate the yellow mini-gene positioned downstream of the UZ mini-gene, activating and repressing it, respectively (not depicted).

Although the *UZ* promoter behaves normally when the *>PRE>* cassette is present, this is not the case when it is absent (Fig. 3C). Instead, beginning shortly after the onset of gastrulation, *>UZ^Wnt^*embryos show a pattern of UZ expression that resembles a super-imposition of the canonical *>PRE>UZ^+^* and Wg patterns. First, these embryos initiate and sustain expression of UZ that is restricted to parasegments 6-12, like the intact *>PRE>UZ^Wnt^* transgene. Hence, as in the case of the *>PRE>UZ^Hmx^* transgene, we infer that the neighboring *Wnt* complex contains a redundant PRE that can substitute for the excised transgene PRE to initially activate the *UZ* promoter in parasegments 6-12 and to maintain the silenced state in the remaining parasegments. Second, the *>UZ^Wnt^* transgene is also strongly up-regulated in all cells that express Wg, seemingly in place of CD2 expression from the *Tub* promoter in the intact *>PRE>UZ^Wnt^* transgene. Accordingly, we infer that the presence of the transgene PRE normally prevents the *UZ* promoter from being co-opted by the *Wnt* complex regulatory elements that are active in these cells, a function that cannot be compensated for by the putative redundant PRE in the Wnt complex. It is possible that the Wnt locus elements act preferentially on the *Tub* promoter to the exclusion of the *UZ* promoter when the *>PRE>* is present but switch to co-opting the *UZ* promoter when the *Tub* promoter is absent. If so, their ability to make this switch does not appear to depend on whether the *UZ* promoter is active or silenced by the redundant PRE in the Wnt complex, as UZ is expressed in Wg-like stripes both in front and behind the anterior boundary of parasegment 6. Alternatively, the combined actions of the transgene and inferred Wnt locus PREs may create a repressive barrier to *UZ* expression that is too high to be breached by the Wnt locus enhancers.

In the thoracic discs, both UZ and CD2 are expressed in *wg*-like patterns (Figs. 3D) indicating that both the *UZ* and *Tub* promoters are now co-opted by Wnt locus enhancers. This contrasts with the embryo, in which the UZ promoter of the intact transgene is stably repressed in cells that will give rise to the wing disc as well all of the more anterior discs (Fig. 3A). Hence, Wnt complex enhancers that operate during larval life can over-ride PRE/PRC dependent silencing of the UZ promoter established and maintained in the embryo.

Importantly, the *wg*-like patterns of UZ and CD2 observed in the discs differ from each other as well as from the native pattern of *wg* in ways that provide insight into the functional interactions between PREs and neighboring enhancer, silencer and promoter elements. We focus on three such differences observed, respectively, in the notum, hinge and wing blade primordia within the wing imaginal disc.

#### (i)#Quantitative rather than qualitative repression of target gene expression in the notum primordium

*wg* is normally expressed in a broad stripe of cells within the prospective notum, the dorso-proximal portion of the wing disc destined to form most of the fuselage of the adult thorax (N; Figs. 3D,4A) and the same is true for the *Tub.CD2* gene within the *>PRE>* cassette. However, UZ is expressed in a much narrower stripe, in cells that express peak levels of Wg and CD2 (Figs. 3D, 4A). Strikingly, *UZ* expression expands to a broad domain resembling that of native Wg and CD2 when the PRE is removed (Figs. 3E,4B). Hence, the transgene PRE reduces—but does not abolish—the response of the *UZ* promoter to the *Wnt* enhancers acting in the notum, indicating that it imposes a quantitative rather than a qualitative barrier to *UZ* expression. We posit that this repressive barrier is sufficiently strong to overcome the activating inputs conferred by the notum *Wnt* enhancer(s) except in cells in which they are maximally active (as revealed by cells that express coincident, peak levels of native Wg and CD2 from the intact transgene, and UZ from the PRE excised *>UZ^Wnt^* transgene).

#### (ii)#Distinct, context-dependent integration of enhancers and PREs in the hinge primordium

*wg* is also expressed in inner and outer rings of cells in the hinge primordium, which surrounds the distal wing primordium (IR and OR in Figs. 3D, 4A). In the IR, CD2 is strongly expressed in contrast to UZ, which is repressed unless the *>PRE>* cassette is excised (Figs. 3D,E and 4A-C). Hence, in this case the relevant Wnt enhancers appear to have co-opted the *Tub* promoter whilst being unable to over-ride silencing of the *UZ* promoter, much as we observe during embryogenesis (Fig. 3B). In the OR, however, the response is the opposite. In this case, UZ is strongly expressed regardless of whether the *>PRE>* cassette is present or absent, whereas the *Tub.CD2* mini-gene in the intact *>PRE>UZ^Wnt^* transgene is repressed. The lack of CD2 expression is consistent with the *Tub* promoter being actively silenced by the transgene PRE, although we cannot test this by excising the PRE as the *Tub.CD2* mini-gene is contained within the *>PRE>* cassette. Unexpectedly, the transgene PRE also appears to repress the native *wg* promoter located ∼ 50 kb away. OR cells that are homozygous for the intact *>PRE>UZ^Wnt^* transgene lack detectable expression of Wg as well as CD2, indicating that the transgene PRE is acting at long range to block transcription of the native *wg* gene (Fig. 3C, top right panel), even though it has no effect on the neighboring UZ promoter. This deficit is associated with the loss of adult structures, in particular, the proximal axillary cord [a hinge structure normally derived from OR cells (Neumann and Cohen, 1996)]. In this case, we can test by excision experiments if the transgene PRE is responsible for silencing the *wg* promoter, and find that this indeed the case: both Wg expression and hinge development are fully restored by PRE excision (Fig. 4C). Thus, the IR and OR rings appear to constitute distinct developmental contexts in which the PRE acts in opposite ways to silence either the *UZ* promoter, or alternatively, the distant *wg* promoter (and possibly the abutting *Tub* promoter).

#### (iii)#Perpetuation of transient gene expression in the wing primordium

*wg* is normally expressed in a thin stripe of border cells flanking the dorso-ventral (D/V) compartment boundary in the prospective wing blade, and the same is true for CD2 from the *Tub.CD2* mini-gene inside the *>PRE>* cassette. In contrast, UZ is expressed inappropriately throughout the prospective wing (Figs. 3D; 4A,C). Moreover, this abnormal expression is eliminated in the absence of the *>PRE>* cassette (Figs. 3E; 4B), as well as cell autonomously when the cassette is excised selectively in the P compartment (Fig. 4C). Thus, within the context of the prospective wing blade, the transgene PRE appears to be sustaining, rather than repressing, *UZ* promoter activity.

Intriguingly, *wg* expression is normally induced early in larval life in most or all cells of the nascent wing primordium by short-range Delta/Serrate signaling across the D/V compartment boundary (Diaz-Benjumea and Cohen, 1995; Doherty et al., 1996; de Celis et al., 1996). However, as the wing grows and descendants of these cells are displaced from the D/V boundary, they stop receiving peak Delta/Serrate signals and cease expressing *wg*. Hence, we suggest that the *Wnt* enhancers that normally initiate *wg* transcription in D/V boundary cells are sufficiently strong to over-ride PRE-mediated silencing of the *UZ* promoter, and that once their descendants stop receiving peak Delta/Serrate input, the transgene PRE acts as a Trithorax Response Element (TRE) to sustain transcription, as has been reported for other genes (Maurange and Paro, 2002; Fujioka et al., 2008; Perez et al., 2011; Bieli et al., 2015). In support, knock-down of the TrxG gene *trithorax* by RNAi causes a similar loss of persistent UZ expression to that resulting from PRE excision (Fig. S1A).

In contrast, expression of the *Tub.CD2* minigene is not perpetuated in descendants of D/V border cells, despite abutting the PRE within the *>PRE>* cassette (Figs. 3D, 4A), just as it is not silenced by the PRE in the canonical *>PRE>UZ^+^* insertion. Hence, it appears refractory to PRE input in both contexts. It is also notable that the transgene PRE acts to reduce expression of both the native *wg* promoter as well as the *UZ* promoter of the intact *>PRE>UZ^Wnt^* transgene in D/V border cells, even as it sustains UZ expression in the descendants of these cells as they proliferate away from the D/V boundary (Fig. 4C).

In sum, the *UZ, Tub* and native *wg* promoters all appear to be integrating both activating and repressing inputs mediated by neighboring PREs, enhancers and other regulatory elements in their vicinity. For the *UZ* and *wg* promoters, the effect of the transgene PRE can be to block, reduce or perpetuate transcription, with the response depending in diverse ways on the developmental history, position and prospective fate of each cell (Fig. 4D).

### Late, long-range, and selective silencing by a transgene PRE inside the continuously expressed *pumilio* gene

The third, atypical insertion, *>PRE>UZ^pum^*, is located inside a large, ∼60kb intron encoding segment of the posterior determinant gene *pumilio* [*pum*; Fig. 5A; (Macdonald, 1992)], and has several properties that provide further insights into the function, range and mechanism of transcriptional repression by PREs.

**Figure 5.**
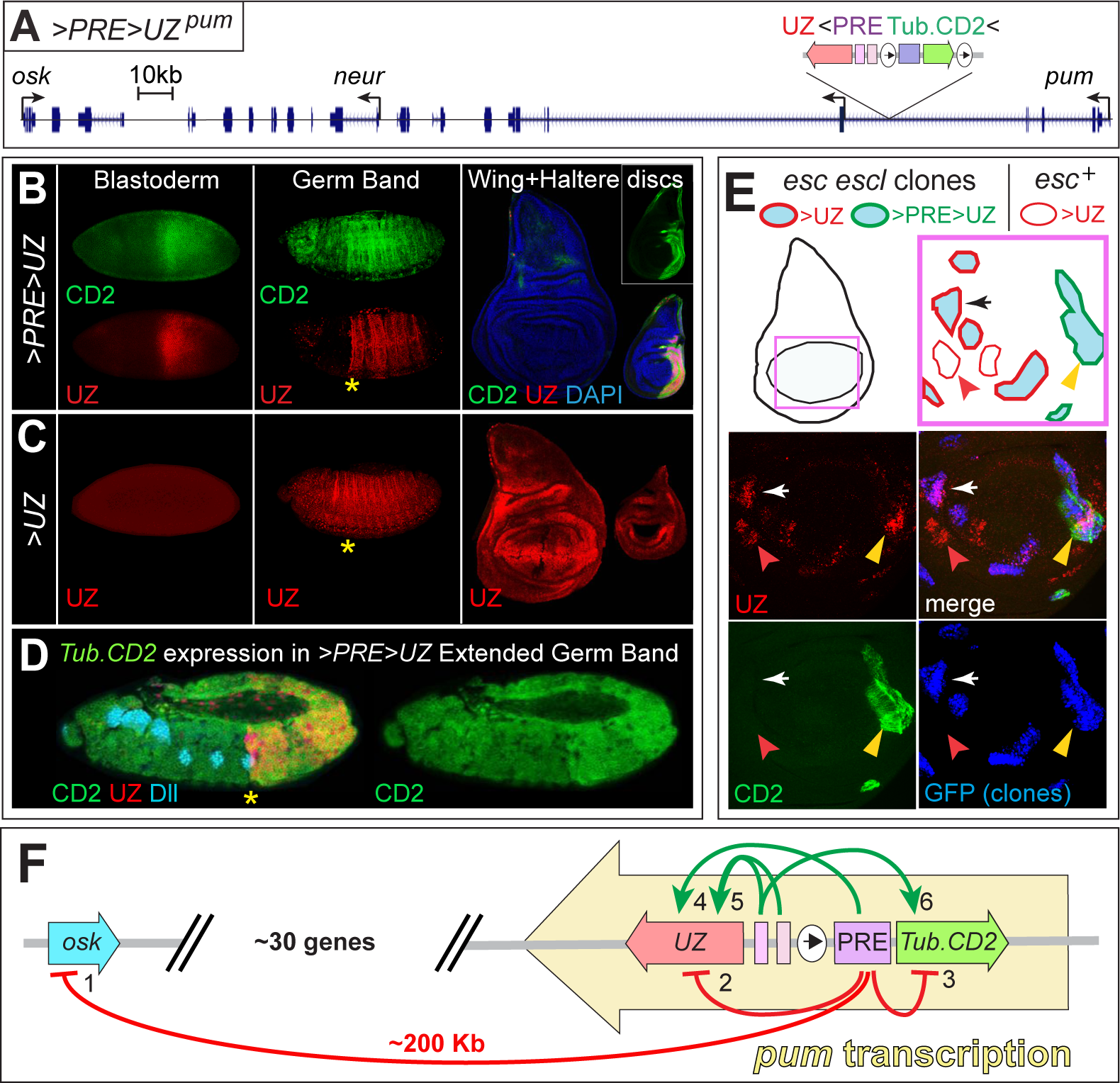
PRE-dependent silencng of the *Tub* promoter in the >PRE> cassette of the *>PRE>UZ^pum^*transgene. **A)** Genomic site and orientation of the *>PRE>UZ^pum^* transgene inside the 160kb intron encoding portion of the *pumilio* locus (transgene not to scale). **B)** Expression of UZ and CD2 in the blastoderm, germ band shortened embryos, and dorsal thoracic discs of *>PRE>UZ^pum^*animals (oriented and annotated as in Figs. 1-3). As in canonical *>PRE>UZ^+^* embryos (Fig. 1A), UZ is activated in parasegments 6-12 in early embryos, and silenced in the remaining parasegments during subsequent embryogenesis, and CD2 is expressed in all cells, albeit up-regulated in parasegments 6-12 in early embryos. Strikingly, and in contrast to the canonical transgene, CD2 expression is subsequently silenced in the wing disc and A compartment of the haltere disc, adopting the same expression pattern as UZ. **C)** Expression of UZ in *>UZ^pum^*embryos and imaginal discs. As in canonical *>UZ^+^* animals, UZ expression is not activated in early embryos but is subsequently expressed ubiquitously, albeit with regional variations in level similar to those observed for the canonical transgene (Fig. 1B). **D)** Expression of UZ and CD2 in a germ band extended *>PRE>UZ^pum^* embryo, counterstained for Dll (turquoise), which marks the nascent primordia of the imaginal discs. CD2 is expressed in all of the embryonic precursor cells that will give rise to the wing and haltere discs, irrespective of whether UZ is silenced or active. In contrast, the transgene PRE mediates the subsequent silencing of *Tub.CD2* expression in the same imaginal discs and compartments in which UZ expression is heritably silenced from the blastoderm stage onwards (**B**). **E)** Silencing of *Tub.CD2* expression of the *>PRE>UZ^pum^* transgenes during larval life depends on PRC2. The top panel depicts a wing disc containing clones of homozygous *esc escl* mutant cells (labeled positively by GFP staining, blue) in the prospective wing (purple box) that carry either the intact or the PRE-excised form of the *>PRE>UZ^pum^*transgene (depicted as in Fig. 2D). The bottom panel shows the independent UZ, CD2 and GFP stains, as well as the merge. An *esc escl* mutant clone carrying the *>PRE>UZ^pum^* transgene is indicated by a yellow arrowhead (note the co-expression of both CD2 and UZ); another clone carrying the *>UZ^pum^* transgene is indicated by a white arrow (expresses UZ, but not CD2). Additionally, one clone of *>UZ^pum^* cells that retains wildtype *esc escl* function is marked by a red arrowhead (does not express GFP or CD2 but does express UZ). **F)** Diagram of tested interactions between the PREs, enhancers and promoters associated with the *>PRE>UZ^pum^*transgene. The transgene PRE silence both the *UZ* (2) and *Tub* (3) promoter, as well as the promoter of the *osk* gene (1) located ∼ 200kb downstream of the UZ mini-gene and separated from it by ∼ 30 protein coding genes (http://flybase.org/). The transgene PRE also acts early in embryogenesis, in conjunction with the EE to activate the *UZ* (4,5) and likely also the *Tub* (6) promoter in parasegments 6-12. The *>PRE>UZ^pum^* transgene resides within a large intron of the *pum* gene, which is expressed throughout development, beginning during the blastoderm stage (yellow arrow); hence the PRE initiates and sustains silencing of both the *UZ* and *Tub* promoters despite the entire transgene being continuously transcribed at a sufficient level for a single copy of the *pum* allele carrying the transgene to sustain wildtype *pum* function.

#### (i)#Late silencing and context-dependent TrxG/PcG regulation of the *Tub* promoter within the *>PRE>* cassette

The *>PRE>UZ^pum^*insert behaves similarly to the canonical *>PRE>UZ^+^* transgene with respect to UZ expression and the requirement for the PRE. The *UZ* promoter is initially activated in parasegments 6-12 and heritably silenced elsewhere in a PRE-dependent fashion (Figs. 5B, compare with Fig. 1A). Likewise, excision of the PRE in the imaginal discs results in a loss of silencing with similar spatial and kinetic parameters [Fig. 5C; as in (Coleman and Struhl, 2017)].

However, with respect to CD2 expression, the *>PRE>UZ^pum^* transgene differs.

Although both *>PRE>UZ^+^*and *>PRE>UZ^pum^* express CD2 ubiquitously during embryogenesis, including in all the founding, progenitor cells of the thoracic discs [marked by expression of the transcription factor Distalless, Dll (Cohen et al., 1993), turquoise; Figs. 5B,D], the behavior of the *>PRE>UZ^pum^* transgene diverges in the imaginal discs, where CD2 expression is now silenced in all cells in which *UZ* is silenced—even though it was not silenced in the embryo (Fig. 5B, right panel). To assess if repression of the *Tub*.*CD2* mini-gene within the *>PRE>* cassette is PRC2 dependent, we applied the same test used above for the *>PRE>UZ^Hmx^* transgene (Fig. 2D), namely, assaying UZ and CD2 expression in *esc escl* mutant clones. As in the case of the *>PRE>UZ^Hmx^*transgene, we find that some *esc^-^ escl^-^* clones in *>PRE>UZ^pum^/+* wing discs ectopically express both CD2 and UZ whereas others express only UZ (Fig. 5E), from which we infer that silencing of the *Tub* promoter, like that of the *UZ* promoter, requires PRC2 activity.

Taken together, these results indicate that the transgene PRE can act in the discs to impose silencing on the *Tub* promoter even though the promoter was previously active in their founding cells in the embryo (Fig. 5F). Curiously, the late imposition of silencing is only observed in the descendants of embryonic cells in which the UZ promoter was not initially activated (Fig. 5B, right panel), raising the possibility it can only occur in the descendants of cells lacking early TrxG activity at the UZ promoter.

Conversely, maintenance of the ON state for both the *Tub* and *UZ* promoters in P compartment cells of the haltere and third leg discs depends on persistent TrxG activity. Specifically, we assayed CD2 and UZ expression in *>PRE>UZ^pum^*haltere discs in which we knocked down *trx* activity in the dorsal compartment, using *ap.Gal4/UAS.trx^RNAi^*.

Under these conditions, both CD2 and UZ expression are concomitantly reduced in the P compartment as is also the case for native Ubx [the latter resulting in an increase in the size of the disc consistent with a haltere-to-wing homeotic transformation (Fig. S1D)]. In contrast, when we performed the same experiment in haltere discs carrying the canonical *>PRE>UZ^+^* transgene, only UZ and native Ubx expression were compromised, whereas CD2 expression remained unaffected (Fig. S1B,C). Hence, the genomic context of the *>PRE>UZ^pum^*transgene has rendered the *Tub* promoter susceptible to the opposing consequences of PcG and TrxG activity in the discs, in contrast to the canonical *>PRE>UZ^+^* transgene, in which the *Tub* promoter is impervious to both. This susceptibility also depends on developmental context, as the *Tub* promoter of the intact *>PRE>UZ^pum^* transgene is refractory to repression in the embryo.

Finally, we asked if the susceptibility of *Tub* promoter to PRE/PRC2-dependent silencing correlates with its state of H3K27 trimethylation. In the case of the canonical *>PRE>UZ^+^* transgene, heritable silencing of the *UZ* promoter in the wing disc is maintained by PRE-dependent, H3K27 trimethylation that extends from the PRE across the *UZ* mini-gene; however, the *Tub.CD2* mini-gene on the other side of the PRE, which is refractory to silencing, remains free of H3K27me3 (Coleman and Struhl, 2017). In contrast, both the *Tub.CD2* and *UZ* mini-genes in the *>PRE>UZ^pum^* transgene are susceptible to silencing, and both are H3K27 trimethylated in the wing disc (Fig. S2).

Hence, the *Tub* promoter can exist in PRE responsive or PRE refractory states depending on genomic and developmental context and correlating, respectively, with whether it is H3K27 trimethylated or not when the *UZ* mini-gene is silenced.

#### <u>(ii)</u> Selective silencing of the *oskar* gene, in *cis* and at long range, by the *>PRE>UZ^pum^*<u>transgene</u>

Consistent with insertion of the *>PRE>UZ^pum^* transgene in the *pum* locus, females that are homozygous for the *>PRE>UZ^pum^* transgene are sterile, laying eggs that give rise to embryos that have a classic posterior determinant phenotype [failure to repress translation of maternal *hunchback* transcripts posteriorly and the consequent failure to form abdominal segments; Fig. 6A(i); (Hulskamp et al., 1989; Struhl, 1989; Wharton and Struhl, 1991)]. However, the transgene does not compromise *pum* function. Instead, several lines of evidence lead to the striking conclusion that it behaves, genetically, as a PRE/PRC2-dependent, recessive allele of a second posterior determinant gene, *oskar* [*osk*; (Lehmann and Nusslein-Vohard, 1986)], which is located ∼200Kb away (Fig. 5A,F).

**Figure 6.**
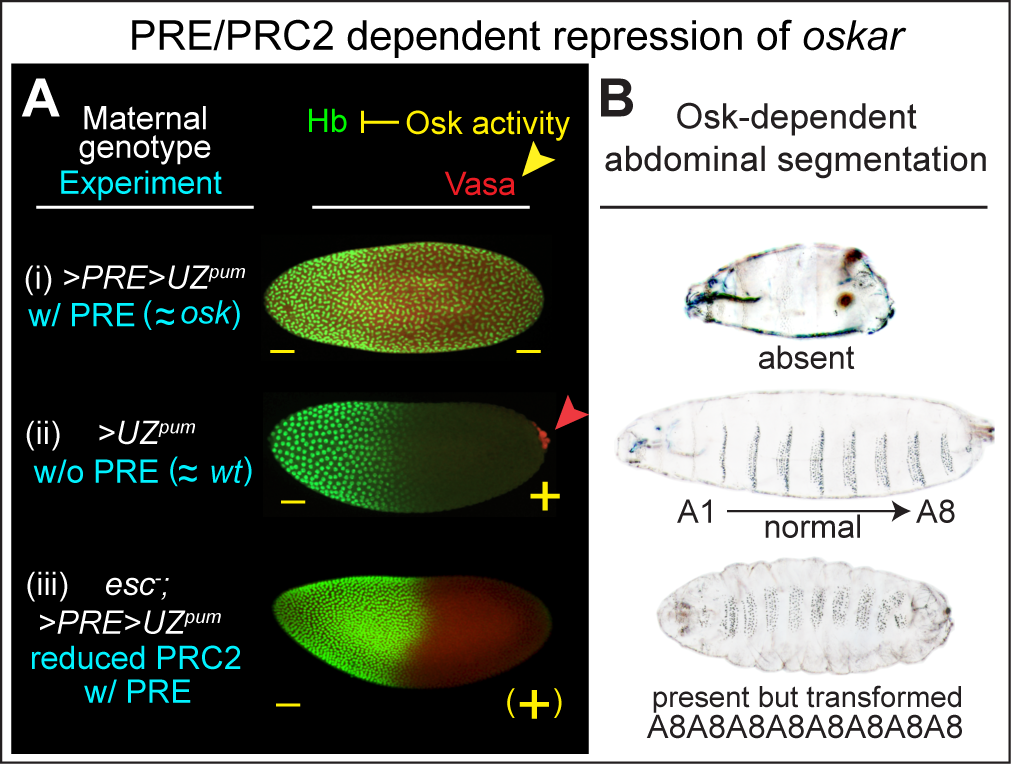
The *>PRE>UZ^pum^* transgene acts in a PRE and PRC2 dependent manner, to silence the *oskar* gene. PRE/PRC2 dependent control of *oskar* (*osk*) activity by the *>PRE>UZ^pum^* transgene as revealed by alterations in Hunchback and Vasa protein expression (**A**) and abdominal segmentation (**B**) caused by deleting the PRE or reducing PRC2 function (maternal genotypes indicated in white; experimental manipulations in turquoise; presence (+), or absence (–) of *osk* activity in yellow). Posteriorly localized maternal *osk* transcripts are required in early embryos for pole cell formation (marked by Vasa protein expression, red) and for suppressing translation of maternal *hunchback* (*hb*) transcripts that would otherwise generate Hb protein (green) and block abdominal segmentation. (i) Embryos derived from females homozygous for the intact *>PRE>UZ^pum^* transgene develop like *osk* null embryos: they lack pole cells, have uniform Hb expression, and lack abdominal segments, indicating that the presence of the transgene PRE has silenced maternal *osk* transcription during the stage it is required to generate localized osk transcripts in the oocyte. (ii) embryos derived from females homozygous for the PRE-excised, *>UZ^pum^*transgene develop like wildtype embryos: they form posterior pole cells, are devoid of Hb protein in the posterior half of the embryo, and form the normal complement of 8 abdominal segments (A1—>A8), indicating that repression of maternal *osk* activity depends on the transgene PRE. (iii) The *osk* null phenotype caused by homozygosity for the *>PRE>UZ^pum^* transgene can be partially rescued by concomitantly removing maternal *esc* function (which greatly reduces, but does not abolish PRC2 activity owing to low level, redundant *escl* gene function). Although pole cell formation is not rescued, Hb protein production is suppressed posteriorly, albeit later than observed for wildtype embryos (by syncytial nuclear cycles 13-14 rather than 9-10) and at least 5 abdominal segments (A1-A5) form, all of which, like all three thoracic segments and at least one head segment, develop as A8-like segments owing to failure to maintain the “off” state of the Bithorax-complex.

First, the *>PRE>UZ^pum^* transgene complements loss of function mutations of *pum* (Experimental Methods). Most incisively, it is possible to maintain a healthy, phenotypically wildtype stock in which the *>PRE>UZ^pum^* transgene is stably balanced by *Df(3R)BSC666*, a deletion that removes the entire *pum* locus. Second, the *>PRE>UZ^pum^* transgene fails to complement loss of function *osk* alleles, including the classic *osk^166^*mutation, as well as the protein and RNA null alleles *osk^A87^* and *Df(3R)osk* (Experimental Methods; Fig. S3). Third, embryos derived from homozygous *>PRE>UZ^pum^* females, as well as transheterozygous *>PRE>UZ^pum^/Df(3R)osk* females fail to segregate pole cells (marked by expression of Vasa protein, red; compare Figs. 6i,ii)—a phenotype specific to the loss of *osk*, but not *pum*, function (Lehmann and Nusslein-Vohard, 1986; Lehmann and Nüsslein-Volhard, 1987). Fourth, excision of the transgene PRE fully restores wild type *osk* gene function (Fig. 6ii); stocks that are homozygous for the *>UZ^pum^* transgene are viable, fertile and phenotypically normal. Fifth, reducing PRC2 function in homozygous *>PRE>UZ^pum^*females by removing *esc* function partially restores the translational repression of *hb* transcripts and promotes abdominal segmentation in their progeny (Fig. 6iii). Thus, the *>PRE>UZ^pum^* transgene acts in a PRE/PRC2-dependent fashion, to repress *osk* gene function during oogenesis.

To determine if the *>PRE>UZ^pum^*transgene acts selectively in *cis* on the *osk* locus, we performed a classic *cis*/*trans* test. Specifically, we compared the capacity of the *pum* allele containing the *>PRE>UZ^pum^* transgene (*pum^>PRE>UZ^*) to suppress *osk* gene activity when located either in *cis* or in *trans* to a sole, wildtype copy of the native *osk* gene. Using the number of pole cells formed in embryos from such “*cis*” or “*trans*” females as a quantitative measure of maternal *osk* gene function, we find that the *pum^>PRE>UZ^* allele acts predominantly, if not exclusively, in *cis*, to repress *osk* gene function (Fig. S3).

The *>PRE>UZ^pum^*transgene is separated from the *osk* locus by around thirty other genes, at least some of which have essential roles during oogenesis, as well as in imaginal discs (e.g., *neuralized*, *tango*; Fig. 5A). Because these functions are not perturbed in homozygous *>PRE>UZ^pum^* animals, it appears that the transgene PRE has acted selectively to silence *osk* without affecting the expression of the intervening genes, consistent with it acting in *cis* to repress *osk* transcription via a locus-selective and long-range looping interaction (Fig. 5F).

#### <u>(iii)</u> Transcriptional elongation through the *>PRE>UZ^pum^* transgene does not compromise <u>either the establishment of silencing or its stability</u>

The location of the *>PRE>UZ^pum^*transgene within an intron-encoding portion of the *pum* gene poses a further question, namely, whether the ability of the PRE to silence the neighboring *Ubx* and *Tub* promoters depends on the absence of read-through *pum* transcription during the establishment and/or maintenance of the silenced state. *pum* is expressed uniformly in all imaginal discs, as verified by antisera against either N- and C-terminal epitopes in the native protein (the specificity of the staining confirmed by loss of signal by RNAi knock-down of *pum*; Fig. 7A, bottom right image; Experimental Methods). Importantly, Pum expression is uniform in both the A and P compartments of haltere discs that are homozygous for the *>PRE>UZ^pum^*transgene, where the *UZ* and *Tub* promoters are silenced in A but active in P (Fig. 7A, lower left panel). We conclude that the transgene PRE has sustained silencing of both promoters during disc development, despite the entire transgene being continuously transcribed within an intron encoding segment of the *pum* gene (and in the case of the *UZ* mini-gene, in the same 5’ to 3’ direction; Fig. 5F).

**Figure 7.**
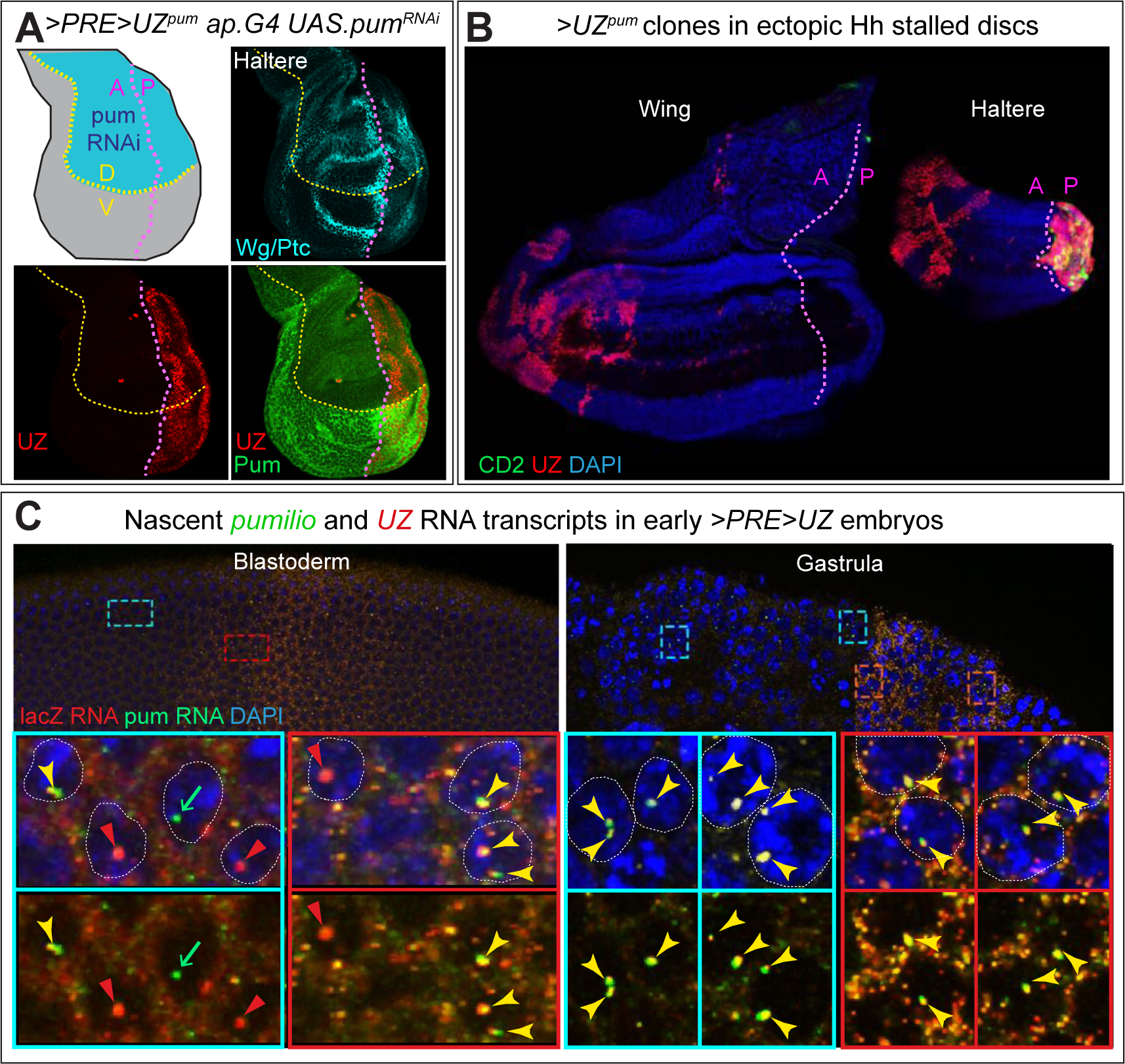
Heritable silencing of the *>PRE>UZ^pum^* transgene embedded within a continuously transcribed intron of *pum*. **A)** Haltere disc homozygous for the *>PRE>UZ^pum^*transgene and expressing RNAi against *pum* transcripts in the dorsal (D) compartment under *ap.Gal4/UAS* control. The D and V compartments are shaded blue and grey, respectively, in the diagram; the A/P and D/V boundaries are indicated, respectively, by magenta and yellow dotted lines, and visualized in Wg and Ptc expression, turquoise). Pum expression, visualized with an antisera against a carboxy-terminal epitope, is greatly reduced in the D compartment validating that the uniform staining observed in the ventral compartment is *bona fide* Pum signal (similar results were obtained with an antisera directed against an amino- terminal epitope). Despite the location of the *>PRE>UZ^pum^* transgene within an intron-encoding portion of the *pum* gene flanked by the coding sequences of the carboxy- and amino- terminal epitopes, UZ expression is “off” in the A compartment and ON in the P compartment, indicating that silencing of the *UZ* promoter in the A compartment is not disturbed by transcriptional elongation through the entirety of the *>PRE>UZ^pum^* transgene and in 5’ to 3’ direction of the *UZ* mini-gene. **B)** Release from silencing following PRE excision from the *>PRE>UZ^pum^* transgene is labile to cell division but refractory to transcriptional elongation. RNAi knock-down of *frataxin* (*fh*) in the larval prothoracic gland under control of the *phantom.Gal4* (*phm.G4*) driver attenuates the normal surge in ecdysone required for pupariation and results in larvae that continue to feed and grow for up to 3 weeks. However, the imaginal discs of such larvae cease growing when they reach their normal, mature size, which occurs 1-2 days after pupation would normally occur and then remain dormant for the remainder of the extended larval life. Growth can be reignited in the A, but not the P, compartments of such “stalled” discs by inducing ectopic expression of Hedgehog (Hh) via the excision of the stop cassette of a *Tub>stop>hh* transgene. *phm.G4 UAS.fh^RNAi^* larvae carrying both the *>PRE>UZ^pum^*and *Tub>stop>hh* transgenes were heat shocked to concomitantly induce *>UZ^pum^* and *Tub>hh* clones around one day before disc growth would otherwise stall, and assayed one week later. Under these conditions, UZ expression, which would otherwise be silenced in the A compartments of both the wing and haltere (Fig. 5B) is now observed in large clone-like A compartment patches that are located far from the A/P compartment boundary and associated with a dramatic expansion in size of the A compartment. However, no such expression is observed in the rest of these discs (aside from normal UZ expression in the P compartment of the haltere disc). Hence, although labile to cell division, silencing of the *UZ* promoter appears stable to continuous transcriptional elongation through the entire *>PRE>UZ^pum^* transgene, even when the PRE is excised. **C)** Ubiquitous transcriptional elongation through the *>PRE>UZ^pum^* transgene in blastoderm (left) and gastrula (right) stage embryos, as detected by fluorescence in situ hybridization (FISH). Regions anterior to, or within, the broad, central stripe of UZ promoter activity in the upper panels that are boxed in turquoise and green, respectively, are shown at higher magnification in the lower panels. The probes used are specific to the *pum* intron in which the transgene is inserted (green) and the *lacZ* coding sequence of the *>PRE>UZ^pum^* transgene (red). The *>PRE>UZ^pum^*transgene is ∼30Kb and embedded in a *pum* intron that is ∼60Kb within a primary transcript of ∼125kb. Hence, nascent transcripts are likely to remain associated with the *pum* locus for at least 45-60 minutes (Shermoen and O’Farrell, 1991) allowing them to be detected as single puncta associated with each allele in each nucleus. Single or double *lacZ* and *pum* puncta are apparent in most nuclei during cellularization of the blastoderm (left panels). At this stage, when transcription has just initiated, the *lacZ* and *pum* signals in the nucleus generally co-localize as is also the case for the gastrula stage, when transcription has reached steady state. Note that cytosolic *lacZ* puncta are also observed in the central region of gastrula stage embryos in which the *UZ* promoter is active, presumably reflecting the accumulation of mature *lacZ* transcripts transcribed under the control of this promoter.

The capacity of the PRE to silence both the *UZ* and *Tub* promoters despite read- through transcription raises the possibility that repression conferred by H3K27 trimethylation of the transgene might be stable to transcriptional elongation. We have previously shown that excision of the PRE from the canonical *>PRE>UZ^+^* transgene results in a progressive, cell-division coupled loss of H3K27 trimethylation and silencing, owing to the reduced read/write capacity of unanchored PRC2 to copy the H3K27me3 mark onto newly incorporated nucleosomes following DNA replication. However, both appear stable indefinitely following excision when cell division is blocked (Coleman and Struhl, 2017). Here, we perform the same test to ask if silencing of the *>PRE>UZ^pum^* transgene is likewise stable in non-dividing cells following PRE excision, despite nucleosome modifications and exchange associated with its being continuously transcribed inside the *pum* intron.

Specifically, we used targeted RNAi mediated knockdown of the gene *frataxin* (*fh*) in the prothoracic gland to generate larvae that cannot pupate. Under these conditions, the imaginal discs cease proliferative growth when they reach normal size, even though the larvae continue to feed and increase in body mass for up to three weeks. Such “stalled” discs can be induced to resume growth in the A, but not the P, compartment by ectopic expression of the secreted protein Hedgehog [Hh; (Coleman and Struhl, 2017)].

As previously observed for the canonical *>PRE>UZ^WT^* transgene (Coleman and Struhl, 2017), inducing ectopic Hh in stalled wing discs carrying the *>PRE>UZ^pum^*transgene causes a dramatic expansion of the A compartment, as well as large clone-like patches of UZ expressing cells in anterior portions of the A compartments. However, no such derepression of UZ is observed in the P compartment nor in more posterior portions of the A compartment where ectopic Hh does not induce new growth (Fig. 7B). Hence, we conclude that silencing of the *UZ^pum^* promoter following PRE excision is labile to cell division but refractory to the level of transcriptional elongation driven by the *pum* promoter over a vastly extended time period.

Although the level of read-through transcription driven by the *pum* promoter does not compromise silencing of the *>PRE>UZ^pum^* transgene, higher levels appear to do so. We generated an intragenic recombinant between the *pum^>PRE>UZ^* allele and a *pum* allele in which *pum* transcription can be boosted under the direct control of UAS regulatory sequences (*pum^UAS^*). We then used the *ap.Gal4* transgene to drive high levels of transcription of the *pum^>PRE>UZ^ pum^UAS^* recombinant allele in the dorsal compartment of the wing disc, beginning midway through the second larval instar, when the *>PRE>UZ^pum^* transgene would otherwise be stably silenced. Under these conditions, the expression of Pum protein (monitored by antisera directed against the amino- and carboxy- terminal epitopes) is elevated 2-3 fold in the D compartment (consistent with a several fold increase in transcriptional elongation through the *>PRE>UZ^pum^*transgene) and silencing is alleviated (Fig. S4).

Finally, we have tested if initiation, as distinct from maintenance, of silencing is refractory to transcriptional elongation by asking if transcription of the *pum^>PRE>UZ^*allele has already begun before the *>PRE>UZ^pum^* transgene is silenced by the PRE. To do so, we assayed for nascent, unspliced *pum^>PRE>UZ^*transcripts by fluorescent *in situ* hybridization (FISH), using probes for *pum* and *lacZ* sequences within the large intron-encoding portion of the *pum^>PRE>UZ^* allele harboring the transgene. Given their exceptionally large size [∼125kb; (Macdonald, 1992)], the primary *pum^>PRE>UZ^* transcripts should remain tethered to the *pum* locus for at least 45-60 minutes, facilitating their detection as single puncta associated with each chromosome (Shermoen and O’Farrell, 1991) and distinguishing such nascent transcripts from mature, maternally deposited *pum* mRNAs and zygotic *UZ* transcripts generated by the *UZ* promoter. Indeed, beginning during cellularization of the blastoderm (nuclear cycle 14), we detect single pairs of fluorescent puncta that stain with either or both the *pum* and *lacZ* probes in most nuclei (Fig. 7C), and such pairs are readily detected in virtually all nuclei as *pum^>PRE>UZ^*transcription rises after the onset of gastrulation (Fig. 7C). Thus, *pum^>PRE>UZ^* transcription begins during the cellular blastoderm stage, but does not compromise the establishment of silencing of the resident *>PRE>UZ^pum^* transgene, which is likely to occur one to two hours after the onset of gastrulation (Struhl and Akam, 1985).

Together, these results indicate that PREs can initiate and maintain heritable silencing of a target gene even when both the PRE and gene promoter are continuously transcribed (Fig. 5F). Moreover, this silencing can be sustained indefinitely by unanchored PRC2 following excision of the PRE, as long as proliferation is blocked to prevent replication coupled dilution of H3K27me3 marked nucleosomes. However, the ability of Gal4/UAS driven over-expression through the *>PRE>UZ^pum^* transgene to overcome silencing suggests that PRC2 anchored at the transgene PRE cannot sustain H3K27 trimethylation against the challenge of several fold higher levels of transcriptional elongation.

## Discussion

### Regulation of heritable versus dynamic modes of transcription by PcG and TrxG complexes

Current views of transcriptional regulation by Polycomb and Trithorax Group complexes remain heavily influenced by the HOX gene paradigm as first elucidated in Drosophila (Yu et al., 2019; Gentile and Kmita, 2020). In this paradigm, HOX genes that are initially repressed are trimethylated at Histone H3K27 by Polycomb Repressive Complex 2.

PRC2 then acts by a “read-write” mechanism to copy the H3K27me3 epigenetic “mark” following DNA replication, and Polycomb Repressive Complex 1 (PRC1) recognizes the mark and acts to impose the OFF state. Conversely, HOX genes that are initially activated are trimethylated at Histone H3K4 and H3K36 by TrxG complexes: this excludes them from H3K27 trimethylation and sustains the ON state. Components of the PcG and TrxG systems also engage in positive and negative feedbacks that reinforce their opposing actions so that the ON and OFF states, once established, are stably propagated from one cell generation to the next. Accordingly, PcG and TrxG complexes are typically viewed as defining alternative, bistable states of chromatin structure and transcriptional regulation, even though these states can be switched under certain conditions.

Strikingly however, both Drosophila and vertebrate PcG and TrxG complexes modify hundreds if not thousands of other target genes, most of which are not locked into stable ON and OFF states. Instead, these genes are regulated in dynamic and quantitative fashion, in response to changing spatial, temporal and/or physiological cues [(Schwartz et al., 2006; Oktaba et al., 2008); reviewed in (Kuroda et al., 2020)]. This poses the general question of how their control by PcG and TrxG complexes relates to the HOX paradigm of heritable epigenetic states.

Here, we address this question by exploiting the requirement for Polycomb Response Elements (PREs) to anchor PcG and TrxG complexes to target genes in Drosophila. This has allowed us to use a transgenic form of a classic HOX gene, *Ultrabithorax.lacZ* that is controlled by a single, excisable PRE *(>PRE>UZ*) to probe how PcGs and TrxGs regulate neighboring genes depending on genomic and developmental context.

Of seven random genomic insertions of the *>PRE>UZ* transgene we have analyzed, four behaved as expected, recapitulating *Ubx*-like patterns of expression throughout development. However, the remaining three, showed distinct and complex deviations from the norm. This is consistent with the prevalence of such position effects as first reported in early studies of PRE-containing transgenes(Simon et al., 1993; Chan et al., 1994; Christen and Bienz, 1994; Sigrist and Pirrotta, 1997; Horard et al., 2000; Poux et al., 2002). Although such “mis-behaved” insertions indicate that the ability of PREs to confer heritable states of expression depends on local genomic milieu, analysis has typically focused on the canonical role of PREs as manifest in “well-behaved” insertions. Here, however, we focus on the anomalous insertions to gain insight into how transcriptional regulation by PRE-recruited PcG and TrxG complexes is integrated with the actions of other transcription factors recruited to the same target genes.

As depicted in the summary diagrams in Figures 2E, 4D and 5F, we have observed a surprisingly diverse collection of PRE-dependent regulatory activities. These include (i) quantitative down-regulation as opposed to a qualitative silencing of transcription, (ii) initiation of heritable gene activity, (iii) repression of a previously active promoter, (iv) inappropriate perpetuation of otherwise transient gene expression, and (v) selective silencing, in *cis* and at long range, of a distant promoter. Significantly, all of these regulatory events appear dependent on genomic and developmental context, presumably conferred by transcription factors associated with neighboring enhancer, silencing and promoter elements.

Accordingly, we posit that the “read-write” capacity of PRC2 to copy H3K27 trimethylation (Margueron et al., 2009; Oksuz et al., 2018), coupled to the capacity of the mark to recruit PRC1 (Cao et al., 2002; Fischle et al., 2003; Min et al., 2003; Wang et al., 2004b), provides the potential for the heritable regulation of gene expression. However, whether this potential is realized depends on the properties of the target genes—in particular whether or not they reside in a permissive genomic milieu, as defined by the absence of cis-acting elements that can counteract the consequences of PcG and TrxG complex activity.

By contrast, we suggest that the many different regulatory interactions we observe for the *>PRE>UZ* insertions in the *Hmx*, *Wnt* and *pum* loci reflect the more general roles of PRE-anchored PcG and TrxG complexes. Specifically, we posit that they control the expression of hundreds if not thousands of target genes scattered throughout the genome by constraining or reinforcing the actions of other transcriptional regulators. For many target genes, this may quantitatively reduce or increase transcription rather than initiate or maintain stable “ON” or “OFF” states of gene expression.

### Context-dependent susceptibility of genes to PRE/PcG/TrxG action

Strikingly, our results provide evidence for a previously unknown plasticity in the control of gene expression by PcG and TrxG complexes—namely the capacity of some target genes to toggle between refractory and responsive states. In principle, whether a given gene is susceptible or impervious to PRE/PcG/TrxG action could be an intrinsic property of the gene, e.g., encoded in sequence-specific elements of its promoter. However, our results monitoring the response of the *Tub.CD2* mini-gene within the *>PRE>* Flp-out cassette argue that this not the case.

Befitting its role in the expression of a “housekeeping” gene, the *Tub* promoter is normally active at moderate level in all cells. In the majority of randomly inserted *>PRE>UZ* transgenes, it appears to be impervious to the presence or absence of the abutting PRE, or to loss of function of either PRC2 or Trx. It is therefore striking that for two transgenes, unique only by their insertion sites in the *Hmx* and *pum* loci, the *Tub* promoter switches from being impervious during embryogenesis to being responsive to both PRC2 and Trx in the imaginal discs. Thus, we infer that the capacity to exist in distinct responsive or refractory states, or to toggle between these states, is conferred by regulatory elements in the neighboring DNA.

### Heritable silencing in the context of read-through transcriptional elongation

Several possible mechanisms have been proposed to explain the opposing actions of PcG and TrxG complexes on gene expression (Schuettengruber et al., 2017; Kuroda et al., 2020; Blackledge and Klose, 2021). Prominent amongst those for mediating silencing by PcG complexes are chromatin compaction and recruitment of H3K27 trimethylated chromatin into nuclear puncta (Polycomb bodies), which reduce or eliminate the accessibility of target genes to nuclear factors required to initiate or sustain transcription (Pirrotta and Li, 2011; Boettiger et al., 2016; Wani et al., 2016; Kundu et al., 2017; Lau et al., 2017; Eeftens et al., 2021). Conversely, there is structural evidence that TrxG complexes countermand silencing by catalyzing H3K4 and H3K36 trimethylation as well as H3K27 acetylation (Papp and Müller, 2006; Tie et al., 2009; Schuettengruber et al., 2009; Tie et al., 2014; Streubel et al., 2018), which preclude trimethylation of H3K27 on the same histone molecule (Yuan et al., 2011; Schmitges et al., 2011; Finogenova et al., 2020). Transcription itself may also oppose silencing by evicting and replacing H3K27me3 nucleosomes with H3.3 nucleosomes (Mito et al., 2007; Deal et al., 2010) and by the direct inhibition of PRC2 enzymatic activity by nascent RNA transcripts (Cifuentes-Rojas et al., 2014; Kaneko et al., 2014; Wang et al., 2017).

These models are all challenged by our demonstration that a *>PRE>UZ* transgene inserted inside an intron encoding segment of the *pum* gene is appropriately and heritably silenced in a PRE-dependent fashion despite being continuously transcribed throughout development. Notably, silencing of the *>PRE>UZ^pum^*insertion is associated with H3K27 trimethylation of the entire transgene and persists indefinitely, even when local anchoring of PRC2 is abolished by PRE excision. This contrasts with the quantal, replication-coupled loss of H3K27 trimethylated nucleosomes and silencing following PRE excision in proliferating cells (Coleman and Struhl, 2017; Laprell et al., 2017). Hence, PRE-anchored PRC2 appears essential to maintain H3K27 trimethylation against the challenge of replication-coupled incorporation of naïve nucleosomes, but not that of continuous transcriptional readthrough—at least at the levels driven by the native *pum* promoter.

Significantly, silencing of the *>PRE>UZ* transgene inserted at the *pum* locus can be overcome by a several fold increase in the level of read-through transcription. This supports a competitive, quantitative relationship between PcG and TrxG complex activities. In Drosophila, such competition could occur at many different levels (Reinig et al., 2020; Holoch et al., 2021), such as the kinetics of compaction, recruitment into repressive nuclear domains, nucleosome exchange or nucleosome modification.

Additional mechanisms, such as DNA methylation, histone demethylation and the divergent activities of multiple, functionally distinct forms of PcG and TrxG complexes, likely contribute in vertebrates (Schuettengruber et al., 2017; Reinig et al., 2020; Kuroda et al., 2020; Blackledge and Klose, 2021). Any one or more of these mechanisms might account for more dynamic and quantitative modes of transcriptional regulation mediated by PcG and TrxG complexes.

### Heritable regulation of HOX genes as a reflection of evolutionary selection for determinants of segmental state

Depending on the criteria applied, the Drosophila genomic contains ∼500-1500 PRE elements interspersed amongst ∼ 20,000 genes (Schwartz et al., 2006; Kwong et al., 2008; Oktaba et al., 2008; Erceg et al., 2017; Bredesen and Rehmsmeier, 2019).

Although many PREs appear to be located within ∼1kb of their presumed target genes, there is ample evidence that they can engage in long range looping interactions between distant genomic sites mediated by PcG proteins, in particular by the oligomerization of components of PRC1 (Lo et al., 2012; Isono et al., 2013; Eagen et al., 2017; Ogiyama et al., 2018; Loubiere et al., 2020). Assuming an average of 1 PRE for every 20 genes, our demonstration that the PRE of the *pum >PRE>UZ* insertion can silence a target gene over 200Kb away, separated by an interval that contains ∼ 30 intervening genes suggests that many if not most genes are located within meaningful, striking range of at least one PRE.

Given the distances over which PREs can act, as well as their prevalence in the genome, we posit that they impose a fundamental constraint on how many, if not most, genes evolve. This may entail the emergence of regulatory elements that render them refractory to PRE/PcG/TrxG action or that allow them to drive the necessary levels and patterns of transcription despite the regulatory impact of PRE-anchored PcG and TrxG complexes. As we observe for the *>PRE>UZ* insertion in the Wnt complex, introducing a single PRE can cause complex quantitative and qualitative changes in transcriptional activity of susceptible target genes that are alleviated by PRE excision, as has also been observed in other contexts (De et al., 2019; Erokhin et al., 2021). Hence, as we have previously argued (Coleman and Struhl, 2017), native PREs scattered throughout the genome have the potential to exert similarly complex and diverse impacts on the transcriptional activities of genes within their realms of influence. Accordingly, we posit that the genomic landscape of regulatory constraints conferred by PREs provides the context in which other *cis*-acting elements controlling gene expression emerge and evolve.

In the case of HOX loci, we argue that the biological imperative for HOX genes to function as determinants of segmental state has rendered them subject to stringent selection against enhancer elements that can over-ride PcG mediated silencing, and in favor of additional PRE elements that would reinforce silencing. As such, we view the heritable control of transcription, whether of HOX or other genes, as an attribute of the target genes and not a fundamental property of the PcG/TrxG machinery upon which this regulation depends.

## Acknowledgements

We thank Atsuko Adachi and Chunyao Tao for technical assistance, Tulsi Patel for determining the insertion sites of the *>PRE>UZ* transgenes, Robin Wharton, Paul Macdonald, Ruth Lehmann and Richard Mann for anti-sera, Vince Pirrotta, Paul Macdonald and Filip Port for the *esc^6^ escl^d01514^* double mutation, the *osk* alleles and transgenes, and the *wg^GFP^* knock-in allele, respectively, and Paul Macdonald, Jürg Müler, Kevin Struhl and Andrew Tomlinson for advice and discussion. We gratefully acknowledge funding from an HHMI Investigatorship and the NIH (R01 GM113000, R35GM127140) to G.S., and pre-doctoral Training grants from the Department of Genetics and Development (5T32GM007088) and the Department of Biochemistry and Molecular Biophysics (T32DK07328) to R.C.

## Experimental Methods

### Constitution and initial characterization of the *>PRE>UZ* transgenes

The canonical *>PRE>UZ* transgene has been reported previously (Coleman and Struhl, 2017). As depicted and described in Fig. 1, it is constructed in the Carnegie 2A P-element vector from previously defined genomic fragments from the *Ubx* gene, as well as the lacZ coding sequence and a *>Tub.CD2>* Flp-out cassette (Jiang and Struhl, 1995) containing the classic 1.6kb *bxd* PRE (see Table 1 for a listing of the sequences defining the ends of each fragment; the full sequence is available on request); the transgene also carries a ∼ 8kb rescuing genomic fragment carrying the *yellow^+^*gene posited downstream of the *UZ* mini-gene as a transgene marker. Seven independent genomic insertions, identified by rescue of the *y* loss of function phenotype, were characterized in detail. Four behaved similarly and showed the expected patterns of UZ and CD2 expression whereas the remaining three showed anomalous patterns of either or both UZ and CD2 as described herein. The sites of insertion were determined by standard inverse PCR (Ochman et al., 1988) and are presented in Table 2.

### Transgenes and mutant alleles

In addition to the *>PRE>UZ^+^*, *>PRE>UZ^Hmx^*, *>PRE>UZ^Wnt^* and *>PRE>UZ^pum^* transgenes (Table 2), the following mutant alleles and transgenes were employed (http://flybase.org/; additional citations below):

*esc^2^*, *esc^6^*, *escl^d01514^*, *wg^GFP^* (Port et al., 2014) *osk^166^*, *osk^A87^*, *Df(3R)osk* (Reveal et al., 2010), *P(osk+)* (Jones and Macdonald, 2015), *P(osk^bcd3-UTR^)* (Ephrussi and Lehmann, 1992), *Df(3R)BSC666*, *P(XP)pum^d04225^/+*, *ap.Gal4*, *hh.Gal4*, *phm.Gal4*, *tub>y^+^>hh* (Basler and Struhl, 1994), *UAS.pum^RNAi^* (Menon et al., 2009), *UAS.fh^RNAi^*(Anderson, 2005), *UAS.trx^RNAi^* (P{TRiP.HMS00580}attP2) and standard *hsp70.flp*, *Tub.Gal80*, *Tub.Gal4*, *UAS.GFP*, *UAS.dcr*, *UAS.flp*, *FRT39*, and *FRT82* transgenes.

#### Exact genotypes (by Figure)

Figs. 1A,B: >PRE>UZ^+^/+ and >UZ^+^/+

Figs. 2B,C: >PRE>UZ^Hmx^/+ and >UZ^Hmx^/+

Fig. 2D: hsp70.flp Tub.Ga4 UAS.GFP; esc^6^ escl^d01514^ FRT40/Tub.Gal80 FRT40;>PRE>UZ^Hm^/+

Figs. 3B-E, 4A,B: >PRE>UZ^Wnt^/wg^GFP^ and >UZ^Wnt^/wg^GFP^

Fig. 4C: >PRE>UZ^Wnt^/>PRE>UZ^Wnt^; hh.Gal4/UAS.flp

Figs. 5B-D: >PRE>UZ^pum^/+ and >UZ^pum^/+

Fig. 5E: hsp70.flp Tub.Ga4 UAS.GFP; esc^6^ escl^d01514^ FRT40/Tub.Gal80 FRT40; >PRE>UZ^pum^/+

Fig. 6 (maternal genotypes): (i) >PRE>UZ^pum^/>PRE>UZ^pum^, (ii) >UZ^pum^ >UZ^pum^, (iii) esc^2^/esc^6^; >PRE>UZ^pum^/>PRE>UZ^pum^, (iv) P(osk^bcd3-UTR^); >PRE>UZ^pum^/>PRE>UZ^pum^, (v) P(osk^bcd3-UTR^)/+; >UZ^pum^ >UZ^pum^.

Fig. 7A: UAS.dcr/+; ap.Gal4/+; >PRE>UZ^pum^ UAS.pum^RNAi^/>PRE>UZ^pum^

Fig. 7B: hsp70.flp; tub>y^+^>hh UASfh^RNAi^/+; >PRE>UZ^pum^/phm.Gal4

Fig. 7C: >PRE>UZ^pum^/>PRE>UZ^pum^ or Df(3R)BSC666

Fig. S1: ap.Gal4/+; >PRE>UZ^pum^ P(XP)pum^d04225^/+

Fig. S1A: ap.Gal4/>PRE>UZ^Wnt^; UAS.trx^RNAi^

Fig. S1B-D: ap.Gal4/+; UAS.trx^RNAi^ alone or in trans to either >PRE>UZ^+^or >UZ

Fig. S2: >PRE>UZ^pum^ (homozygous or in trans to Df(3R)BSC666) Fig. S3: as indicated

Fig. S4: ap.Gal4/+; >PRE>UZ^pum^ P(XP)pum^d04225^/+

### Chromatin Immunoprecipitation

All ChIP experiments were performed on wing discs from wandering, late third instar larvae and were conducted in 2-3 independent biological replicates. Immunoprecipitation and qPCR analysis were performed as previously described (Coleman and Struhl, 2017). Probes used for qPCR are detail in Table 3.

**Table 3.**
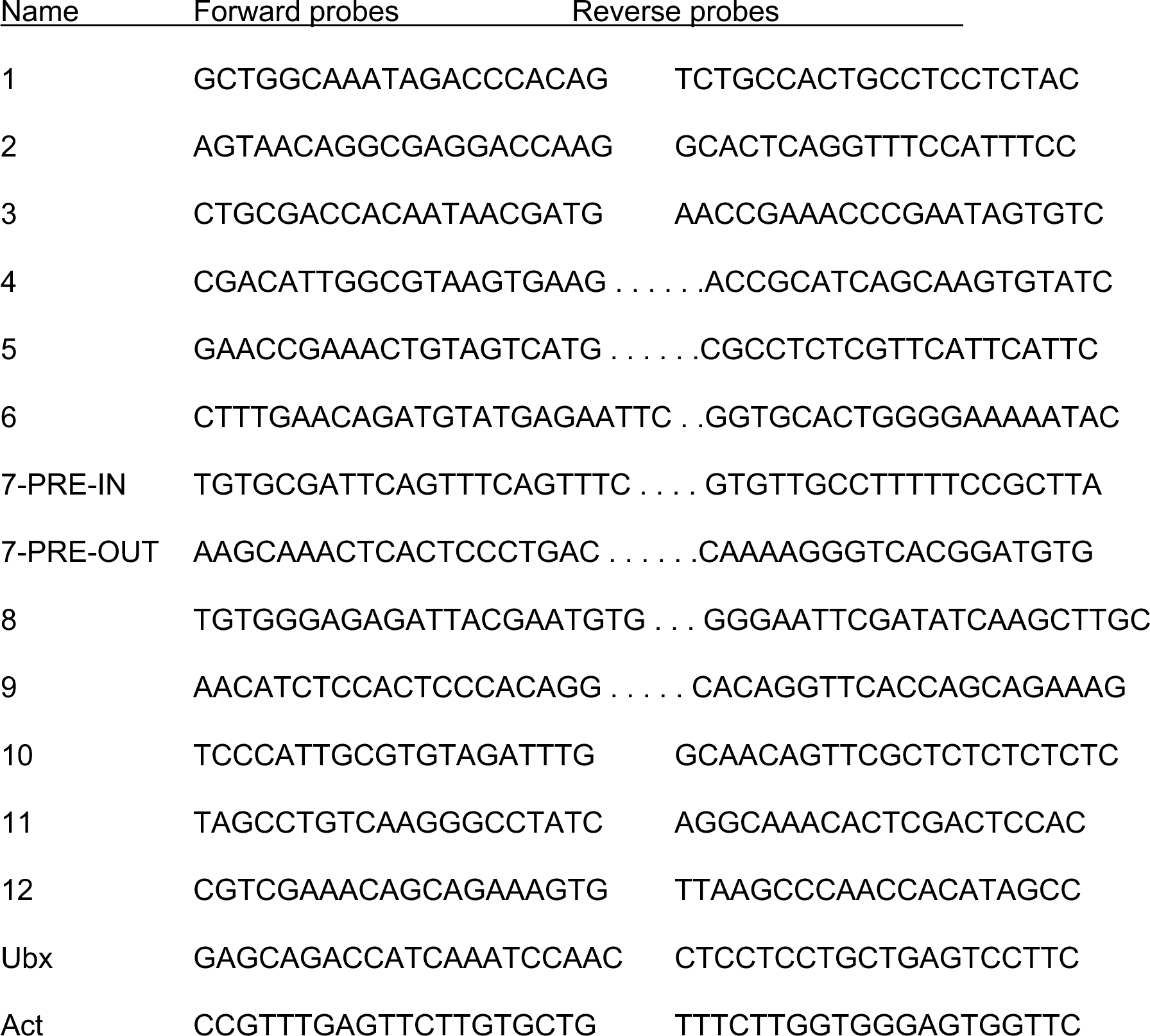
Probes (5’ to 3’) used for qPCR analysis (Fig. S2)

#### Generation of Flp-mediated excision clones with or without positively marked *esc escl* mutant clones

i. *>PRE>* excision clones were generated either by *hh.Gal4* driven expression of a *UAS.flp* transgene (Fig. 4C) or by heat shock induced expression of an *hsp70.flp* transgene [Fig. 7B; in this case coinduced with *Tub>hh* clones ∼1 day before *phm.Gal4 UAS.fh^RNAi^* induced cessation of wing disc growth; (Coleman and Struhl, 2017)].
ii. *esc escl* mutant clones, marked positively by GFP (Figs. 2D, 5E) were generated using the MARCM technique (Lee and Luo, 1999) in genetic backgrounds carrying the desired *>PRE>UZ* transgenes (see exact genotypes).

#### RNAi knock down

i. Dorsal compartment specific knock-down of Pum in haltere discs homozygous for the *>PRE>UZ^pum^* transgene (Fig. 7A) was achieved using *ap.Gal4* to drive expression of *UAS.pum^RNAi^*.
ii. Dorsal compartment specific knock-down of trx in wing discs homozygous for the *>PRE>UZ^Wnt^* (Fig. S1A) or in *wildytpe*, *>PRE>UZ^+^*/+ or *>PRE>UZ^pum^/+* haltere discs (Figs. S1B-D) was achieved using *ap.Gal4* to drive expression of *UAS.trx^RNAi^*

#### Immunofluorescence, in situ hybridization, and microscopy

Protein expression in embryos and imaginal discs were assayed in situ by standard fixation methods and immunofluorescence, using rabbit polyclonal anti-βGal (Cappel, 1/20,000), mouse monoclonal anti-CD2 (BD Pharmingen#OX-34, 1/1000), mouse anti-Wg (DSHB#4D4, 1/30), mouse anti-Ubx (DSHB#FP3.38, 1/50), rat anti-Hb (gift of Paul MacDonald, 1/4000), rabbit amino- and rat- carboxy-termini of Pum (gift of Robin Wharton; 1/1000), guinea pig anti-Dll (gift of Richard Mann; 1/6000), rat anti-Vasa (gift of Ruth Lehmann; 1/1000), and appropriate AlexaFluor 488, 555 and 647 conjugated secondary antisera (ThermoFisher; 1/5000) plus DAPI (Hoechst 33342, Invitrogen) to counterstain to label nuclei. Preparations were imaged using an SP5 Leica confocal microscope.

RNA expression in embryos was assayed by standard methods of in situ hybridization using FISH probes (Kosman et al., 2004). Plasmids for *in vitro* transcription were generated by cloning PCR amplified *pum* and lacZ DNAs into BamHI and NotI or EcoRI and NotI sites, respectively, of pBluescript SK(-).

*pum* Fwd primer: AAAATAGGATCCGCGGCACACCCCAAAAATACAAGAG.

*pum* Rev primer: TATTAAAAAATGCGGCCGCACTCCACCGACCGACTGACTGACAC. lacZ Fwd primer: AAAAAAGAATTCACCCAACTTAATCGCCTTGCAGCAC.

lacZ Rev primer: ATATTTGGATCCAAGGCGGTCGGGATAGTTTTCTTGC.

*In vitro* transcription was carried out with DIG or Biotin-labeled nucleotides (Roche) and hybridized transcripts were stained using sheep anti-DIG (Roche#1333089) or mouse anti-Biotin (Roche#1297597) primary antisera followed by appropriate AlexaFluor488, 555 conjugated secondary antisera. *pum* transcripts from the *pum* allele containing the *>PRE>UZ^pum^* transgene were assayed in homozygous *>PRE>UZ^pum^* embryos derived from females hemizygous for the transgene (in *trans* to a deletion of the *pum* locus, *Df(3R)BSC666*) mated with homozygous *>PRE>UZ^pum^*males (homozygous embryos were distinguished from hemizygous embryos by the presence of two *lacZ*/*pum* RNA staining puncta per nucleus). Sense probes and *in situ* hybridization carried out in embryos that did not carry the transgene served as negative controls to confirm that nuclear signal reflected bona fide nascent transcripts.

---

**Figure S1.**
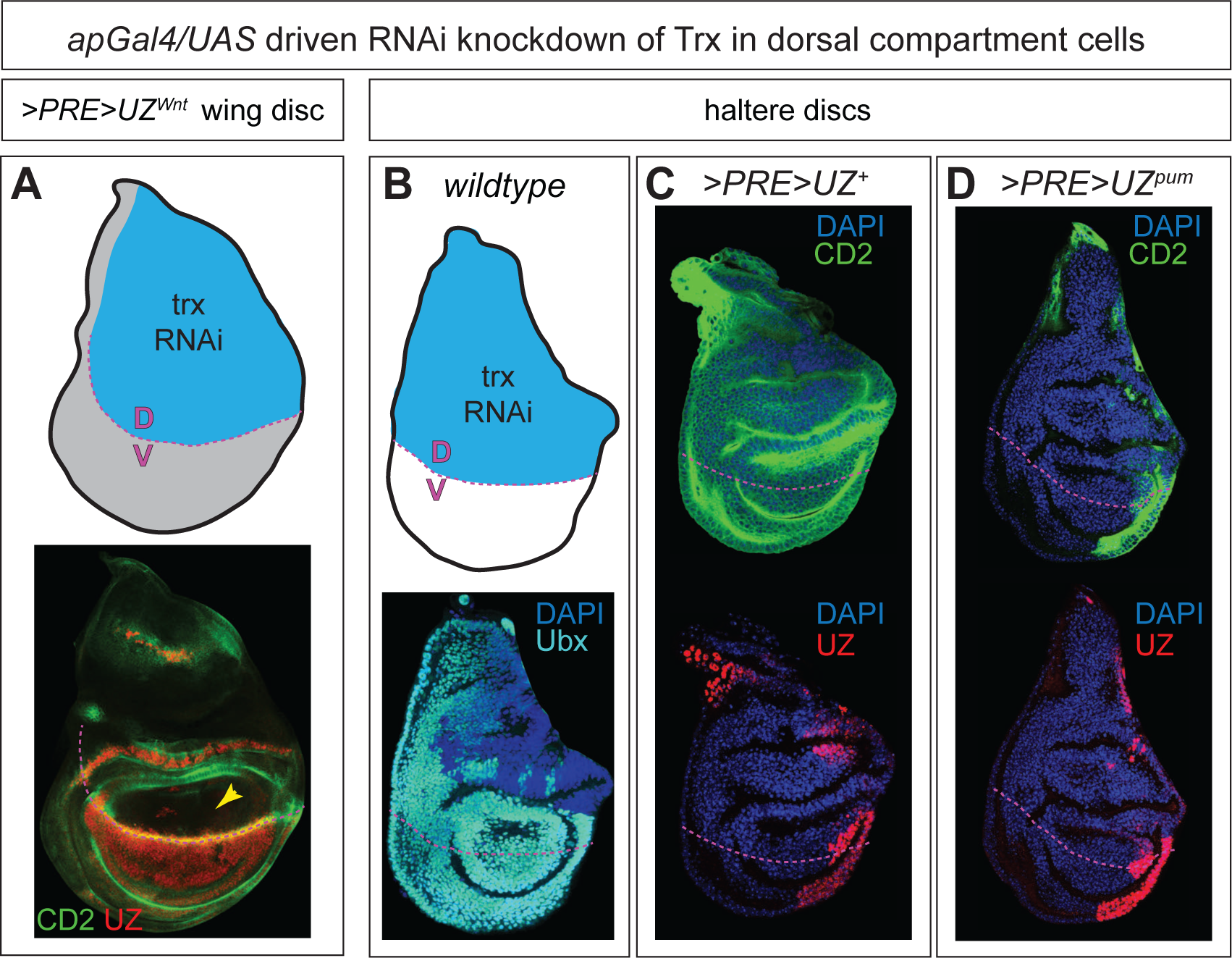
trx is required for persistent *>PRE>UZ^Wnt^* and *>PRE>UZ^pum^* expression in the prospective wing and haltere, respectively. **A)** RNAi knock-down of trx in the dorsal (D) compartment of a *>PRE>UZ^Wnt^*wing imaginal disc (blue in the cartoon) stained for both UZ (red) and CD2 (green) expression. The PRE-dependent expression of UZ throughout the prospective wing (Figs. 3C,D,4A,B) is selectively lost in the D compartment. **B-D)** RNAi knock-down of trx in the D compartments of wildtype (**B**), *>PRE>UZ^+^* (**C**) and *>PRE>UZ^pum^* (**D**) haltere discs. The wildtype disc is stained for Ubx (turquoise) and shows the partial failure to sustain activity of native Ubx promoter. Both UZ and CD2 expression is similarly diminished for the *>PRE>UZ^pum^* transgene (**D**) whereas only the UZ expression is reduced for the *>PRE>UZ^+^* transgene (**C**), indicating that the *Tub* promoter within the *>PRE>* is refractory to TrxG activity in the canonical *>PRE>UZ^+^* transgene, but dependent on TrxG activity in the *>PRE>UZ^pum^*.

**Figure S2.**
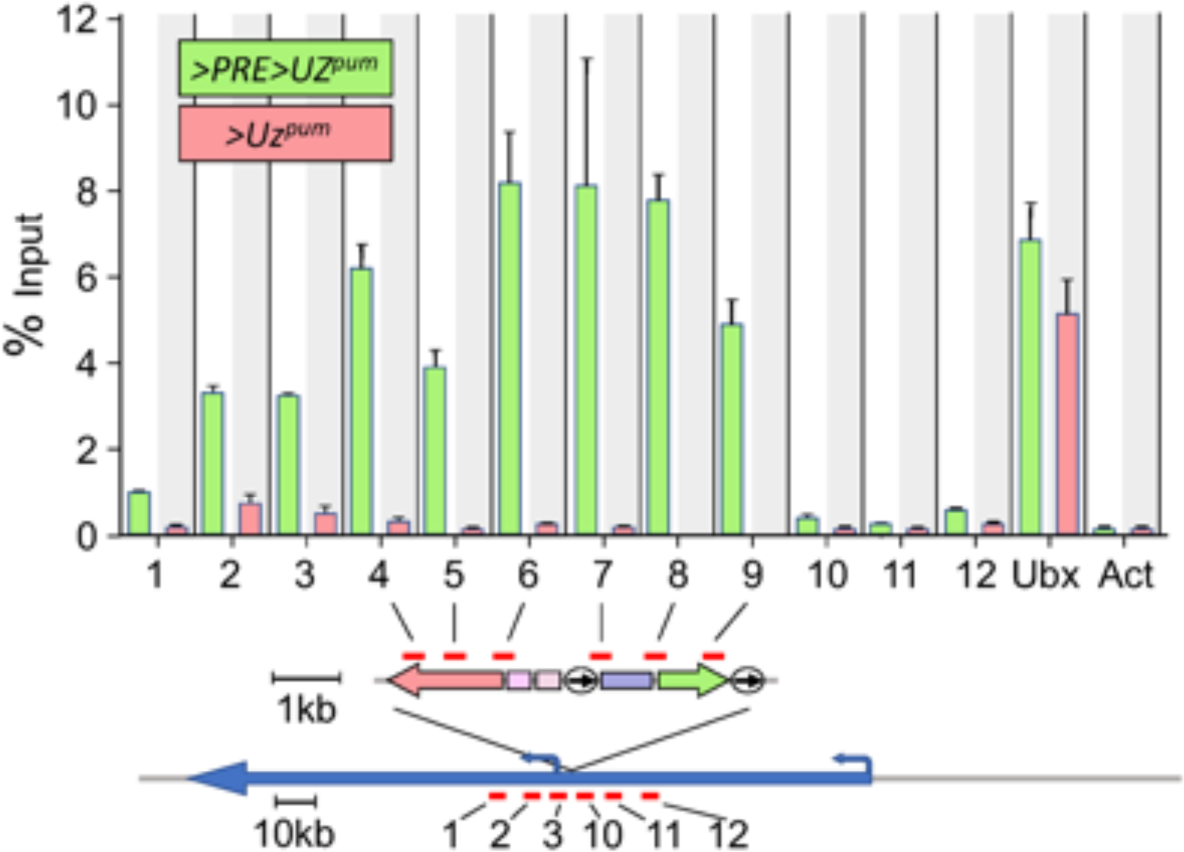
PRE-dependent H3K27 trimethylation of the *>PRE>UZ^pum^* transgene in wing discs. ChIP-qPCR for H3K27me3 for *>PRE>UZ^pum^*(green) and *>UZ^pum^* (red) wing discs. H3K27me3 is detected for all probed regions of the *>PRE>UZ^pum^* transgene (probes 4-9) as well as the downstream portion of the encompassing *pum* gene (probes 1-3). In contrast, no significant H3K27me3 signal was observed for any of probed regions for the *>UZ^pum^*transgene or the adjoining pum genomic DNA (probes 1-12). The native *Ubx* gene (repressed and H3K27 trimethylated in wing discs) and the *Actin5c* gene (expressed and not trimethylated) serve as controls (Papp and Müller, 2006). The corresponding ChIP-qPCR for H3K27me3 for the canonical *>PRE>UZ^+^*and *>UZ^+^* transgenes is similar except that probe 9 shows no detectable H3K27me3 (Coleman and Struhl, 2017). Bars represent the mean ±SEM of 2-3 independent biological replicates. Bars are absent for probes 8 and 9 in the *>UZ^pum^* condition because probed DNA is not present in animals of this genotype. Probes for ChIP-qPCR analysis are presented in Table 3. Note that alternative “PRE-IN” and “PRE-OUT” versions of probe 7 are used to detect the probed fragment in the *>PRE>UZ^pum^* and *>UZ^pum^* forms of the transgene (Coleman and Struhl, 2017).

**Figure S3.**
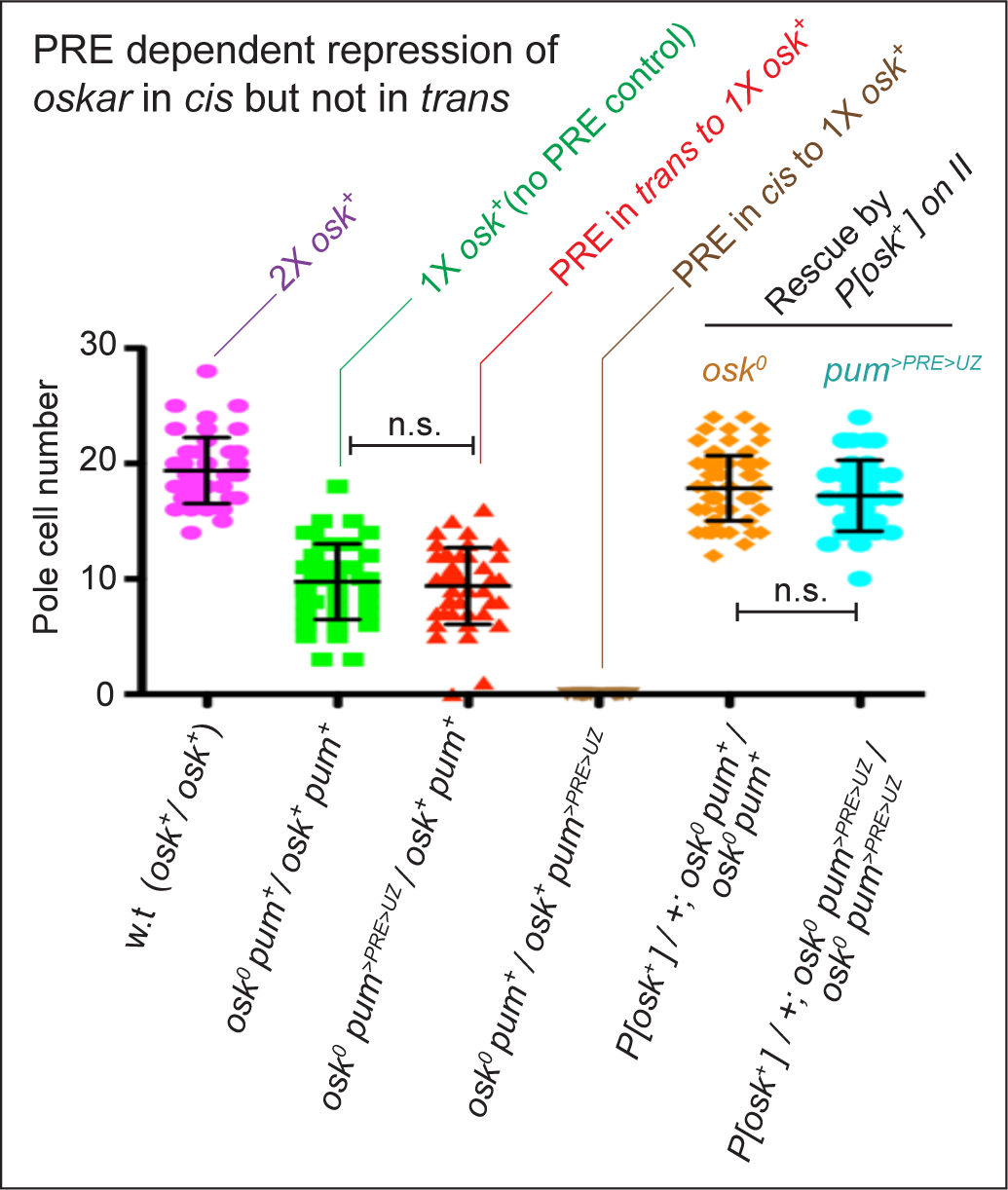
*Cis/trans* test of silencing of the wildtype allele of *osk* by the >PRE>UZ^pum^ transgene. Pole cell number (mean ±SD) in embryos was used as a quantitative measure of maternal *osk* gene activity as validated by a two-fold reduction in pole cell number observed comparing embryos from *w.t.* females (magenta dot plot) versus females heterozygous for a null allele of *osk* (*osk^0^*; green dot plot); P<0.001). Pole cell numbers were not further reduced when the *>PRE>UZ^pum^* transgene was present in *trans* rather than in *cis* to the wildtype *osk* allele (red dot plot). In contrast, when the *>PRE>UZ^pum^* transgene is present in *cis*, no pole cells were observed (brown dot plot; P<0.001). Equivalent results were obtained for embryos from homozygous *osk^0^* females carrying an *osk^+^* rescuing transgene on chromosome II. In this case, homozygosity for the *>PRE>UZ^pum^* transgene in *cis* to the *osk^0^* allele (turquoise dot plot) had no effect on the capacity of the *osk^+^* rescuing transgene to rescue pole formation (orange dot plot), again arguing that suppression of *osk* transcription by the *>PRE>UZ^pum^* transgene, even when homozygous, occurs only in *cis*. As in the text, the *>PRE>UZ^pum^* transgene in these experiments is designated as a *pum* allele, *pum^>PRE>UZ^*, and the *w.t.* and mutant alleles of both the *osk* and *pum* loci indicated in all of the genotypes. Pole cells were visualized by morphology and Vas protein staining and pole cell counts were conducted by an observer blind to the experimental genotype. P values were calculated by unpaired, two-tailed t tests. *w.t.* n=48; *osk^0^ pum^+^/osk^+^ pum^+^* n=42; *osk^0^ pum^>PRE>UZ^/osk^+^ pum^+^* n =44; *osk^0^ pum^+^/osk^+^ pum^>PRE>UZ^* n=20; *P[osk^+^]/+; osk^0^ pum^+^/osk^0^ pum^+^* n=88; *P[osk^+^]/+; osk^0^ _pum_>PRE>UZ*_/*osk*_*0 _pum_>PRE>UZ* _n=43._

**Figure S4.**
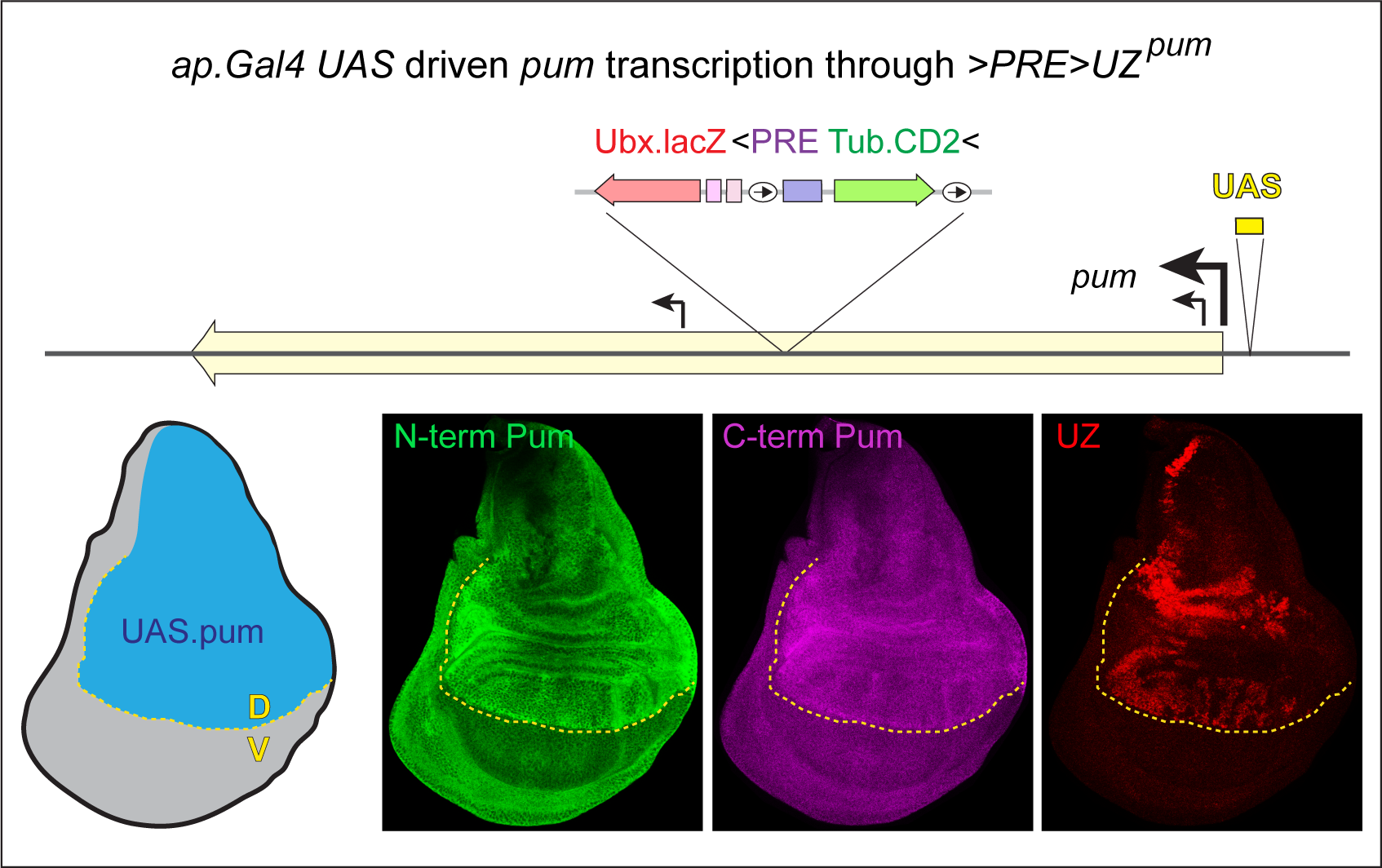
Gal4/UAS activation of the *pum* promoter over-rides silencing of the *>PRE>UZ^pum^* transgene. Wing disc heterozygous for the *pum^>PRE>UZ^ pum^UAS^* recombinant allele carrying the *>PRE>UZ^pum^* transgene downstream of a UAS regulatory element inserted just upstream of the *pum* promoter (top) and expressing Gal4 in the dorsal (D) compartment under the control of the *ap.Gal4* driver. *ap.Gal4* is first expressed during the second larval instar, well after the *>PRE>UZ^pum^*transgene is silenced by the resident PRE during embryogenesis. The D and V compartments in the mature disc are shaded blue and grey, respectively, in the diagram, and the D/V boundary is indicated by the yellow dotted line. Pum expression, visualized with separate antisera against amino- and carboxy-terminal epitopes, is elevated ∼2-3 fold in the D compartment, confirming a several fold increase in transcriptional activity of the *pum* promoter and resulting in a loss of UZ silencing.

